# Coupling of ATPase activity, microtubule binding and mechanics in the dynein motor domain

**DOI:** 10.1101/309179

**Authors:** Stefan Niekamp, Nicolas Coudray, Nan Zhang, Ronald D. Vale, Gira Bhabha

## Abstract

The movement of a molecular motor protein along a cytoskeletal track requires communication between enzymatic, polymer-binding, and mechanical elements. Such communication is particularly complex and not well understood in the dynein motor, an ATPase that is comprised of a ring of six AAA domains, a large mechanical element (linker) spanning over the ring, and a microtubule-binding domain (MTBD) that is separated from the AAA ring by a ~135 Å coiled-coil stalk. We identified mutations in the stalk that disrupt directional motion, have microtubule-independent hyperactive ATPase activity, and nucleotide-independent low affinity for microtubules. Cryo-electron microscopy structures of a mutant that uncouples ATPase activity from directional movement reveal that nucleotide-dependent conformational changes occur normally in one half of the AAA ring, but are disrupted in the other half. The large-scale linker conformational change observed in the wild-type protein is also inhibited, revealing that this conformational change is not required for ATP hydrolysis. These results demonstrate an essential role of the stalk in regulating motor activity and coupling conformational changes across the two halves of the AAA ring.

## Introduction

Dyneins are minus-end directed, microtubule-based molecular motors that belong to the AAA+ (ATPases associated with diverse cellular activities) superfamily of proteins. Cytoplasmic dynein is responsible for the transport of numerous cargoes along microtubules (MTs), such as organelles, vesicles, viruses, and mRNAs (Vale, 2003; Vallee *et al*, 2004). In addition, cytoplasmic dynein plays key roles in facilitating basic cell biological processes such as spindle positioning during mitosis (Kiyomitsu & Cheeseman, 2013). Mutations and defects in cytoplasmic dyneins are associated with many diseases such as neurodegenerative diseases and cancers (Roberts *et al*, 2013).

The cytoplasmic dynein holoenzyme is composed of two identical ~500 kDa heavy chains and multiple associated polypeptide chains that primarily bind to the N-terminal tail of dynein (Pfister *et al*, 2006). Regulatory proteins such as Lis1 and NudE bind to some dyneins and can modify its motility properties (Vallee *et al*, 2012; Kardon & Vale, 2009). To initiate processive motility for cargo transport, human cytoplasmic dynein also requires dynactin as well as cargo-adaptor proteins such as BicD and Hook3 (McKenney *et al*, 2014; Schlager *et al*, 2014). However, the core element for motility of all dyneins lies in the conserved motor domain of the heavy chain, which consists of six different AAA domains that are linked together as an asymmetric hexameric ring (AAA1-AAA6). Only AAA1-AAA4 can bind nucleotides (Burgess *et al*, 2003; Carter *et al*, 2011; Schmidt *et al*, 2015; Kon *et al*, 2012; Schmidt *et al*, 2012; Cho *et al*, 2008; Kon *et al*, 2004) (Fig. 1A); ATP hydrolysis in AAA1 is required for dynein stepping and AAA3 acts as a switch that facilitates robust motility when ADP is bound (Bhabha *et al*, 2016, 2014; DeWitt *et al*, 2015). The catalytic domains in the AAA ring are spatially distant from the microtubule binding domain (MTBD); the two are connected via the coiled-coil “stalk” that emerges from AAA4. Another coiled-coil element, called the buttress, protrudes from AAA5 and interacts with the stalk close to the ring (Fig. 1A). The buttress also has been shown to be important for the allosteric communication between ring and MTBD (Kon *et al*, 2012). The N-terminal linker, which lies on top of the ring, is believed to serve as a mechanical element that drives motility (Burgess *et al*, 2003). Over the last few years, several structural studies have illuminated a series of conformational changes in the dynein AAA ring during the ATPase cycle (Bhabha *et al*, 2014; Schmidt *et al*, 2015; Kon *et al*, 2012; Carter *et al*, 2011). The key conformational changes include domain rotations within the AAA ring and rearrangements of the linker domain.

**Figure 1.**
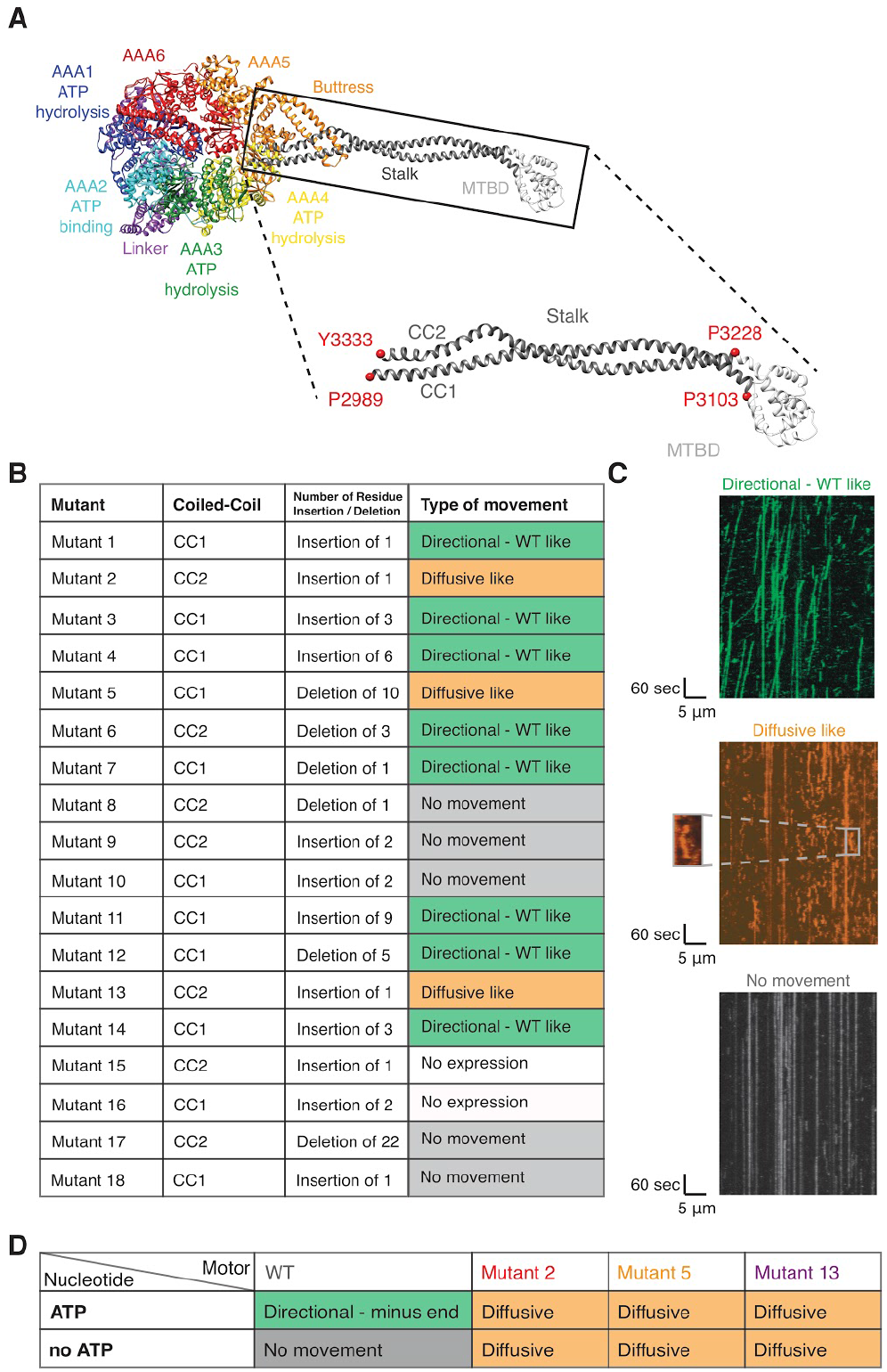
Single-molecule motility properties of dynein stalk mutants reveal mutants with nucleotide-independent diffusive motility. (A) Structure and domain organization of the motor domain of cytoplasmic dynein (PDB 4rh7 (Schmidt et al, 2015)). Inset shows zoom of MTBD (white) and coiled-coil stalk (grey), which consist of two helices, CC1 and CC2. Well conserved residues, that were used as anchor points to define CC1 and CC2 in this study are depicted as red spheres. Numbering is based on yeast cytoplasmic dynein. (B) Table showing location, number of inserted or deleted residues and motility phenotype of all 18 stalk mutants. Examples of single-molecule assay results are shown in Supplementary Fig. 4C-E. Sequence information and exact position of individual mutants are shown in Supplementary Fig 2, 3. Quantification and classification is based on three technical repetitions. (C) Example kymographs for ‘Directional - WT like’ (top), ‘Diffusive like’ (middle), and ‘No movement’ (bottom). Magnified area for ‘Diffusive like’ motion shows run of a single molecule. The kymographs for each mutant are shown in Supplementary Fig. 4C. (D) Table showing the type of movement found for wild-type, mutant 2, mutant 5, and mutant 13 in a modified single-molecule assay with and without ATP. Classification of type of movement is based on two repetitions of different dynein preparations. Kymographs for each mutant are shown in Supplementary Fig. 5. Supplementary Movies 1-4 and 5-8 show motility of wild-type and diffusive mutants with and without ATP, respectively.

To coordinate motility, motor proteins must communicate between the ATPase and polymer binding site. ATP binding to AAA1 results in a weakened affinity (K_d_ >10 *μ*M) of dynein for microtubules (MTs). After ATP hydrolysis, the motor binds MTs with stronger affinity (K_d_ <1 *μ*M) (Kon *et al*, 2009). In this manner, the AAA ring controls the affinity of the MTBD for MTs. Conversely, interaction of the MTBD with MTs regulates the ATPase activity in the AAA ring (Kon *et al*, 2009). How this allosteric communication occurs is still poorly understood. In the case of kinesin and myosin, the ATPase and track binding sites are located relatively close (within ~25 Å) to each other in the same domain (Vale & Milligan, 2000). In dynein, however, the very small ~10 kDa microtubule-binding domain is spatially separated from the AAA ring by the ~135 Å long coiled-coil stalk (Imamula *et al*, 2007; Gibbons *et al*, 2005; Redwine *et al*, 2012; Carter *et al*, 2008; Kon *et al*, 2009). Furthermore, the stalk is positioned between AAA4 and AAA5, which is on the opposite side of the ring from AAA1, resulting in a ~240 Å separation between the main catalytic site and the MTBD.

To enable two-way communication between the MTBD and AAA ring, it has been suggested that the stalk undergoes conformational changes (Oiwa & Sakakibara, 2005; Shima *et al*, 2006; Kon *et al*, 2009). One hypothesis is that sliding between the two antiparallel helices of the stalk coiled-coil leads to changes in their register with respect to each other, with each registry corresponding to different microtubule affinities; the stalk in the β+ registry results in a low MT affinity state and the α registry results in high MT affinity (Kon *et al*, 2009; Gibbons *et al*, 2005). This is further supported by structural work which has shown that when ADP-vanadate (ADP-vi) is bound to AAA1, the coiled-coil 2 (CC2) of the stalk is kinked and slides together with the buttress relative to coiled-coil 1 (CC1) (Schmidt *et al*, 2015). Another study speculates that local melting of the coiled-coil between different states of the hydrolysis cycle plays a major role in the communication (Nishikawa *et al*, 2016; Gee & Vallee, 1998). However, how relative length changes of the stalk either via sliding or local melting drive the communication between the ring to the MTBD is not well understood.

To gain better insights into the allosteric communication between the AAA ring and the MTBD, we have identified mutants in the dynein stalk that block communication between the ATPase and microtubule binding sites. These mutants show diffusive movement along MTs and also hydrolyze ATP at maximal rates in a microtubule-independent manner. Structural characterization by cryo-electron microscopy (cryo-EM) of one of these mutants reveals a stabilization of a previously uncharacterized open conformation of the AAA ring in the presence of the non-hydrolysable ATP analogue AMPPNP. In the presence of ADP-vanadate (ADP-vi), mimicking the post-hydrolysis state of dynein, we observed that this mutant is primed for hydrolysis, but with the linker in an extended conformation, which differs from the bent conformation of the linker in wild-type dynein (Schmidt *et al*, 2015; Bhabha *et al*, 2014). This result reveals that linker bending is not essential for ATP hydrolysis. Moreover, we gained new insights into domain movements in the AAA ring. The cryo-EM structure of the mutant in AMPPNP and ADP-vi states show that one half of the AAA ring undergoes a conformational change similar to the wild-type enzyme, while the AAA domain movements in the other half of the ring, from which the stalk extends, are disrupted. This result reveals that the stalk likely plays a key role in coupling conformational changes throughout the AAA ring. Our results provide insight into how conformational changes are coordinated within dynein’s motor domain to allow microtubule regulation of ATPase activity and motility.

## Results

### Stalk mutants show nucleotide-independent diffusion

Given the spatial separation between dynein’s catalytic AAA ring and the MTBD, it is apparent that allosteric communication must be mediated in some way via the stalk (Fig. 1A). To understand what regions of the stalk may play a role in the allosteric communication, we aligned and analyzed 534 sequences of the dynein motor domain. We found that the length of the stalk is very wellconserved (99% of the sequences have the exact same stalk length) among species and types of dynein, such as cytoplasmic, axonemal and IFT dynein, but the sequence is not (Supplementary Note 1, Supplementary Fig. 1). Based on the conserved length of the stalk and our sequence analyses, we decided to investigate how insertions and deletions in the stalk affect dynein’s motility. We designed a panel of 18 insertion and deletion mutants in the yeast cytoplasmic dynein background, based on our sequence analysis (Fig. 1B, Supplementary Fig. 2, 3, Supplementary Table 1). We expressed and purified GST-dimerized versions of each mutant (Supplementary Fig. 4A) with an N-terminal GFP (Reck-Peterson *et al*, 2006; DeWitt *et al*, 2012), assessed the quality of the protein using negative stain electron microscopy to ensure structural integrity, and used single-molecule total internal reflection fluorescence (TIRF) microscopy assays (Reck-Peterson *et al*, 2006; Yildiz & Vale, 2015) for initial characterization of single-molecule motility. Our panel of mutants displayed a wide variety of phenotypes (Fig. 1B, C). In several locations, insertions and deletions could be tolerated without any effect on motility (Fig. 1B, Supplementary Fig. 4B, Supplementary Note 2). Of these eight mutants that show wild-type phenotypes, seven are in CC1, suggesting that this helix is more tolerant of length changes than CC2 (Supplementary Fig. 1L). One region that is particularly sensitive to mutation is at the interface of the stalk and buttress, which often resulted in a dead motor (Fig. 1B, Supplementary Fig. 1L). This interface has been shown previously to play a role in nucleotide-dependent conformational change (Schmidt *et al*, 2015). Three mutants from our panel (mutants 2, 5, and 13) presented an interesting diffusive-like behavior, with single molecules randomly moving back-and-forth along the microtubule (Fig. 1B, C, Supplementary Fig. 5, Supplementary Movies 1-4). This observation suggests that the motors are weakly bound to microtubules but unable to undergo effective unidirectional motion.

To assess the nucleotide-dependence of the diffusive phenotypes, we carried out single-molecule experiments in the absence of ATP. As expected, the wild-type control showed no movement, and was rigor bound to microtubules. Surprisingly, in the absence of ATP, all three mutants displayed diffusive behavior very similar to that observed in the presence of ATP. Diffusion, that we observed even in absence of nucleotide, suggests that mutant 2, mutant 5, and mutant 13 have a weakened interaction with microtubules (Fig. 1B, Supplementary Fig. 5, Supplementary Movies 5-8).

We also assessed the nucleotide-dependent binding affinity of dynein for microtubules using a cosedimentation assay. In wild-type dynein, the motor binds tightly to microtubules in the absence of ATP, but weakly in the presence of ATP (Fig. 2A). In contrast to the nucleotide-dependent microtubule-affinity of wild-type enzyme, the microtubule affinity of the mutants was low in the absence of nucleotide and in the presence of ATP or AMPPNP (Fig. 2B-D, Supplementary Table 2), which is consistent with the single-molecule motility results. These results confirm that the diffusive mutants have a weakened microtubule affinity which remained unchanged in different nucleotide states.

**Figure 2.**
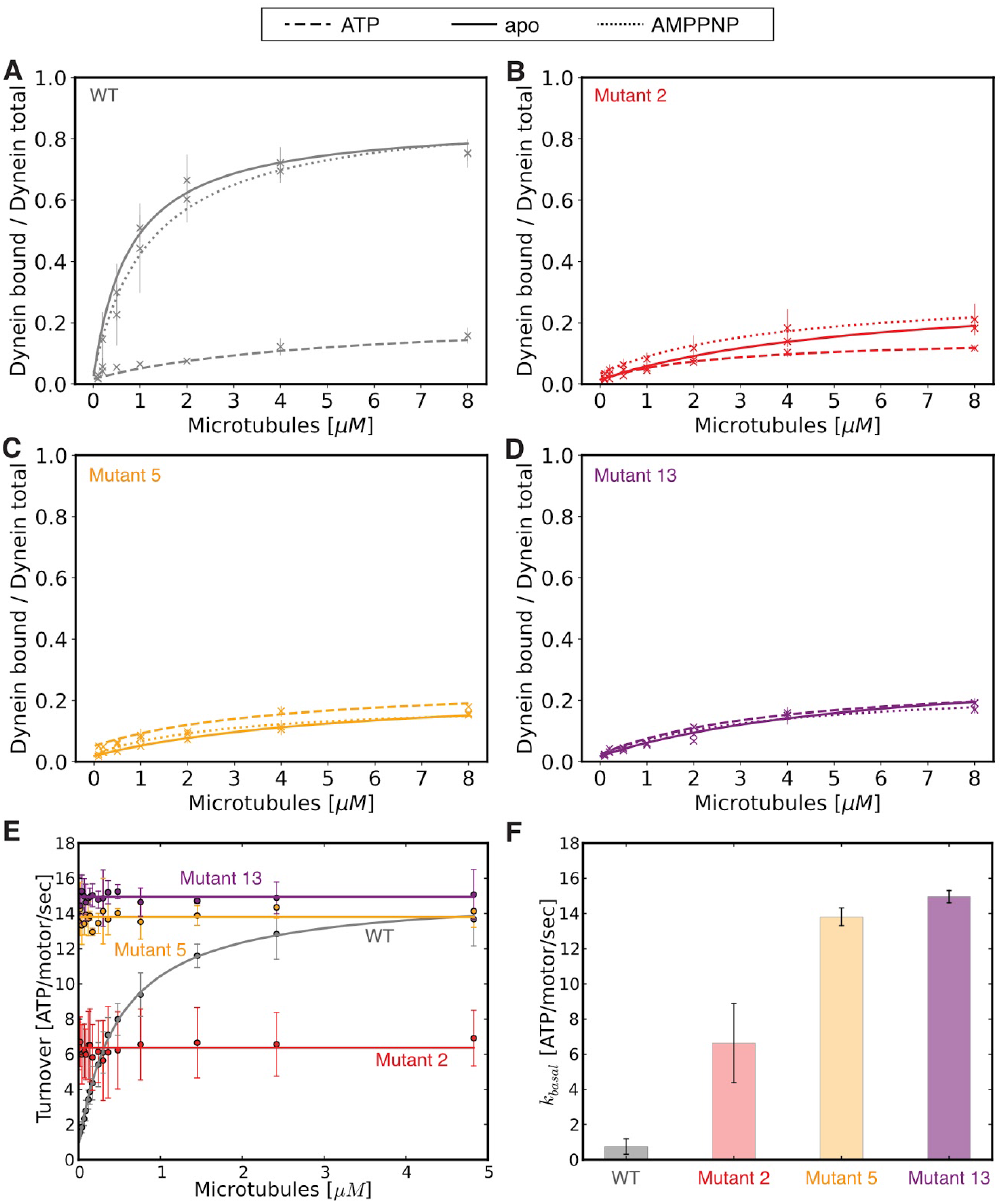
Diffusive mutants show microtubule independent, high basal ATPase activity and low affinity for microtubules. Microtubule affinity measured by a cosedimentation assay in the apo state (full line) and in the presence of ATP (dashed line), and AMPNP (dotted line) for (A) wild-type, (B) mutant 2, (C) mutant 5, and (D) mutant 13. (E) Microtubule stimulated ATPase activity of wild-type (grey), mutant 2 (red), mutant 5 (orange), and mutant 13 (purple). (F) Bar plot of basal ATPase activity of wild-type (grey), mutant 2 (red), mutant 5 (orange), and mutant 13 (purple). Error bars in A-F show standard deviation of three repetitions of different dynein preparations. Supplementary Table 2 and 4 show fit equation and rate quantification for microtubule affinity and ATPase data, respectively.

Since we did not observe any directional movement of these three mutants in single-molecule assays, we asked whether there is any net directionality in a microtubule gliding assay when there are many motors interacting with a microtubule. In this microtubule gliding assay, monomeric dyneins (wild-type or mutants) were attached to a glass coverslip (Supplementary Fig. 6A). Results from this assay show that the three mutants generated microtubule gliding across the glass surface, although their velocities were ~10-fold lower than wild-type dynein (Supplementary Fig. 6B, Supplementary Movies 9-12). In order to determine whether the microtubules were moving in the same direction as for wild-type dynein, we assessed the direction of motion with single molecules of a human homodimeric kinesin-1 (K490) (Tomishige *et al*, 2006a), which move processively towards the plus end (Supplementary Fig. 6A). By observing the direction of kinesin movement along the gliding microtubules, we could assess their polarity. Our results showed that the direction of mutant 2, 5, and 13 in microtubule gliding assays was the same as for wild-type dynein.

In conclusion, mutants 2, 5, and 13 show nucleotide-independent diffusive movement as single molecules, while ensembles of these motors can produce extremely slow directional movement towards the MT minus end.

### Diffusive mutants show microtubule independent hyperactive ATP hydrolysis

Since single-molecule analysis of mutants 2, 5, and 13 showed diffusive movement uncoupled from the nucleotide state, we asked whether these mutants were capable of hydrolyzing ATP. One possible hypothesis was that the mutants could no longer bind or hydrolyze ATP, while another possibility is that ATP hydrolysis was uncoupled from directional movement. We measured the ATPase activity of mutant 2, mutant 5, and mutant 13 at varying concentrations of microtubules. In wild-type dynein, ATPase activity is stimulated in the presence of microtubules, resulting in a characteristic increase in ATPase activity as microtubule concentration increased, until maximal ATPase activity is reached. For wild-type dynein, we measured a basal ATPase turnover of 0.75 ± 0.34 ATP/motor/sec, which increased with increasing concentrations of microtubules to a k_cat_ of 15.18 ± 1.18 ATP/motor/sec and K_m_ of 0.50 ± 0.17 *μ*M for tubulin (Fig. 2E). These ATPase values are similar to those previously reported (Cho *et al*, 2008; Carter *et al*, 2008; Toropova *et al*, 2014) (Supplementary Table 3). Surprisingly, and in contrast to wild-type dynein, the three diffusive mutants showed high basal ATPase activity that did not significantly increase upon the addition of microtubules. Interestingly, the basal ATPase activities of mutants 5 and 13 were very similar to the maximal microtubule-stimulated ATPase activity of the wild-type protein (Fig. 2E, F, Supplementary Table 4). Together, these results indicate that the diffusive mutants hydrolyze ATP in a microtubule-independent manner.

### Structural basis for hyperactivity of mutant 5

Our functional and biochemical assays showed that insertions and deletions in mutants 2, 5, and 13 result in 1) diffusive movement of single dynein molecules on microtubules, 2) constitutively hyperactive ATPase, and 3) constitutively weak microtubule binding that is not modulated by nucleotide. Taken together, these results suggest that these mutations disrupt the two-way communication between the MTBD and AAA ring in the dynein motor domain. Next, we sought to understand the structural basis underlying the uncoupling between the microtubule and ATPase sites in the diffusive mutants. Because of its high basal ATPase activity, we decided to focus on mutant 5.

We first collected a cryo-EM dataset of mutant 5 in the presence of 2 mM AMPPNP to mimic an ATP-bound state of the enzyme at AAA1 and AAA3. After 3D classification and refinement, we identified two distinct classes, with reconstructions at ~7.5-8 Å resolution (Supplementary Fig. 7A, B). This resolution allowed us to establish conformational changes at the subdomain level and model helices in some parts of the structure (Supplementary Fig. 7C). Each AAA domain consists of a large subdomain (AAAL) and a small subdomain (AAAs), which can be considered as rigid bodies in the context of our resolution. Each AAAL and AAAs subdomain is fit independently as rigid bodies into each density map to generate a model corresponding to each map.

The most evident change in the motor domain of the majority (~71% of all particles - class 1, 7.7 Å resolution) of mutant 5 particles was an substantial opening between the small and large domains of AAA5 (Fig. 3A, B), which was previously only observed as a minor conformation for the wild-type motor (Supplementary Note 3, Supplementary Fig. 8). In addition, density for most of the distal stalk as well as the buttress is missing, suggesting that these regions are flexible. For the minor conformation (~29% of all particles - class 2, 7.6 Å resolution), the cryo-EM map shows a closed ring with no gap between the small and large domain of AAA5 and the helices of the initial part of the stalk and for the buttress are well defined (Fig. 3A, Supplementary Fig. 7C). In this class, we can identify a conformation that has previously been referred to as the high microtubule affinity state in which the coiled-coil 2 of the stalk is not kinked (Supplementary Fig. 7C) (Schmidt *et al*, 2015). An additional and more subtle difference between the class 1 and 2 density is found in the N-terminal GFP tag at the end of the linker. In contrast to class 2, for class 1 (major class with “open” ring), the density for the N-terminal GFP tag is not well defined (Supplementary Fig. 7D, E), which may indicate that the N-terminus of the linker is more flexible and potentially undocked from the ring at AAA5. Looking at domain movements in both class 1 and class 2 (Fig. 3C, D), we also found that AAA2L is positioned away from the active site of AAA1 (Supplementary Fig. 7F-R). Since the gap between AAA1 and AAA2 must close for productive ATP hydrolysis, we concluded that the ring of mutant 5 in the AMPPNP state is not primed for hydrolysis, as is true for wild-type dynein.

**Figure 3.**
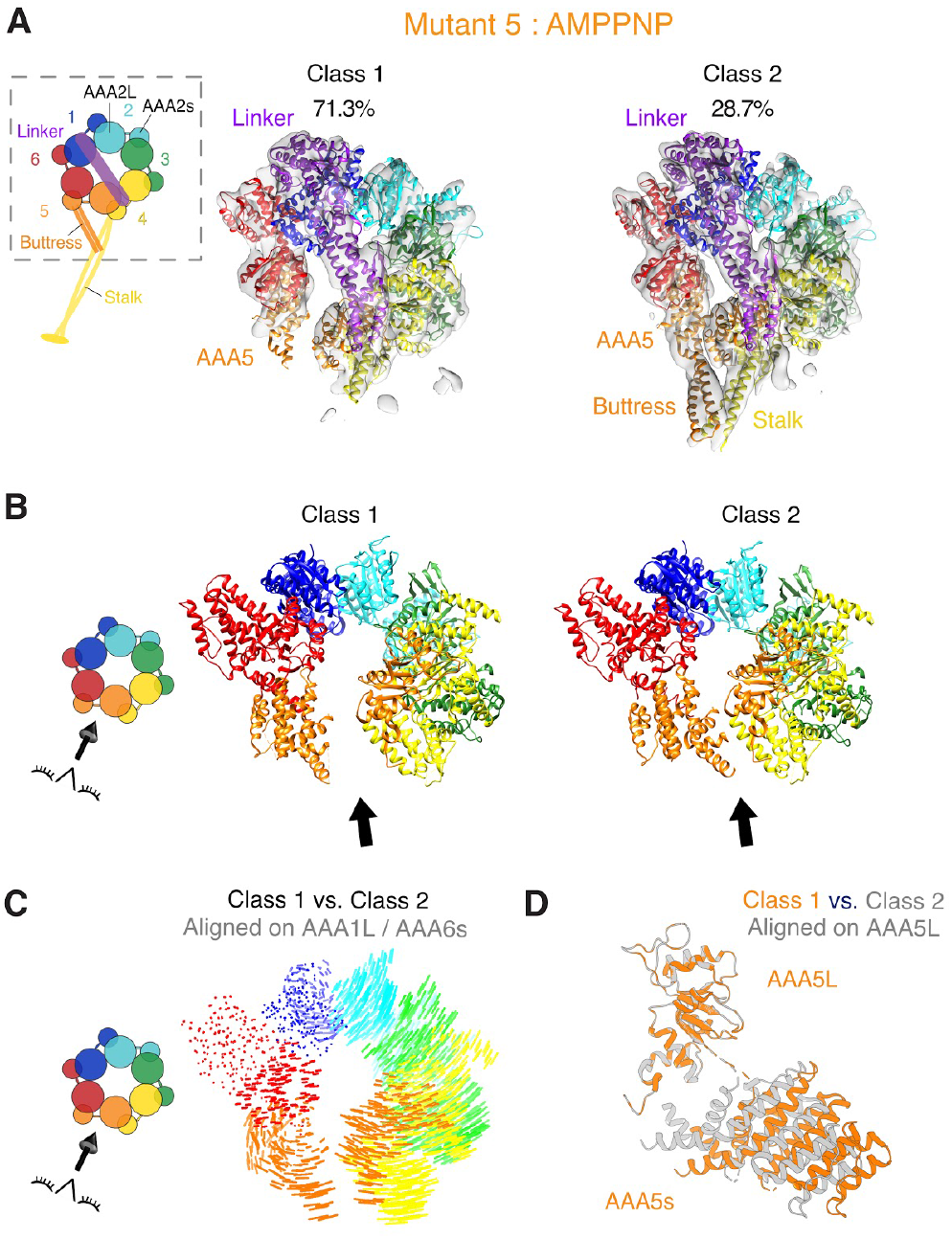
Cryo-EM structure of mutant 5 in the presence of AMPPNP shows a gap in the AAA ring. (A) Cryo-EM reconstructions and fitted models for class 1 and class 2 resulting from 3D classification of the data. Class 1 is composed of 71.3% of all particles (left) and class 2 of 28.7% of all particles (right). The cryo-EM density map for both classes is shown as a semi-transparent surface with a fitted model (fit as described in Materials and Methods) as cartoon. Color coding of domains is the same as for Fig. 1. Left: schematic of monomeric dynein construct, box indicates region that was resolved in the cryo-EM maps. (B) Cartoon representation of models for both classes. Black arrow indicates the position of the gap between AAA5L and AAA5s in class 1. Left: schematic indicates the point-of-view. (C) Visualization of inter alpha carbon distances between class 1 and class 2 as shown in B after alignment on AAA1L. We removed the linker for clarity. Left: schematic indicates the point-of-view. (D) Movements between the large and small domains of AAA5 between class 1 (orange) and class 2 (grey). The large domain of AAA5 is aligned.

We next examined mutant 5 in the ADP-vanadate (ADP-vi) state, which mimics the post-hydrolysis state of dynein (Schmidt *et al*, 2015). In this state, the AAA domains in wild-type dynein adopt a more compact conformation in which the gap between AAA1 and AAA2 closes, which primes AAA1 for nucleotide hydrolysis (Supplementary Fig. 7R). In addition, the linker changes from a “straight” conformation (extended linker spanning from AAA1 to AAA5) to a “bent” conformation (the N-terminus of the linker making contacts with AAA 3/2). Our cryo-EM data for mutant 5 in the presence of 2 mM ATP and 2 mM vanadate resulted in a ~9 Å reconstruction, for which subdomain movements could be mapped with confidence (Fig. 4A, Supplementary Fig. 9A). Based on fitting AAAs and AAAL domains into our density as described above, our data show that the gap between AAA1 and AAA2 closes (Fig. 4B, Supplementary Movie 13), as observed for the human cytoplasmic dynein 2 crystal structure (PDB 4RH7 (Schmidt *et al*, 2015) and the yeast dynein cryo-EM reconstruction (PDB 4W8F (Bhabha et al. 2014)). Unlike the bent linker observed with ADP-vi (Schmidt *et al*, 2015), the mutant 5 linker is not bent at the hinge-point (Fig. 4A). However, the N-terminal region of the linker is undefined, suggesting increased flexibility at its N-terminal region (Fig. 4A, C, Supplementary Fig. 9B). Thus, our structural data for mutant 5 in the ADP-vanadate state indicate that the motor is primed for hydrolysis, but does not undergo the large conformational change in the linker that is believed to be essential for motility (Burgess *et al*, 2003; Bhabha *et al*, 2016).

**Figure 4.**
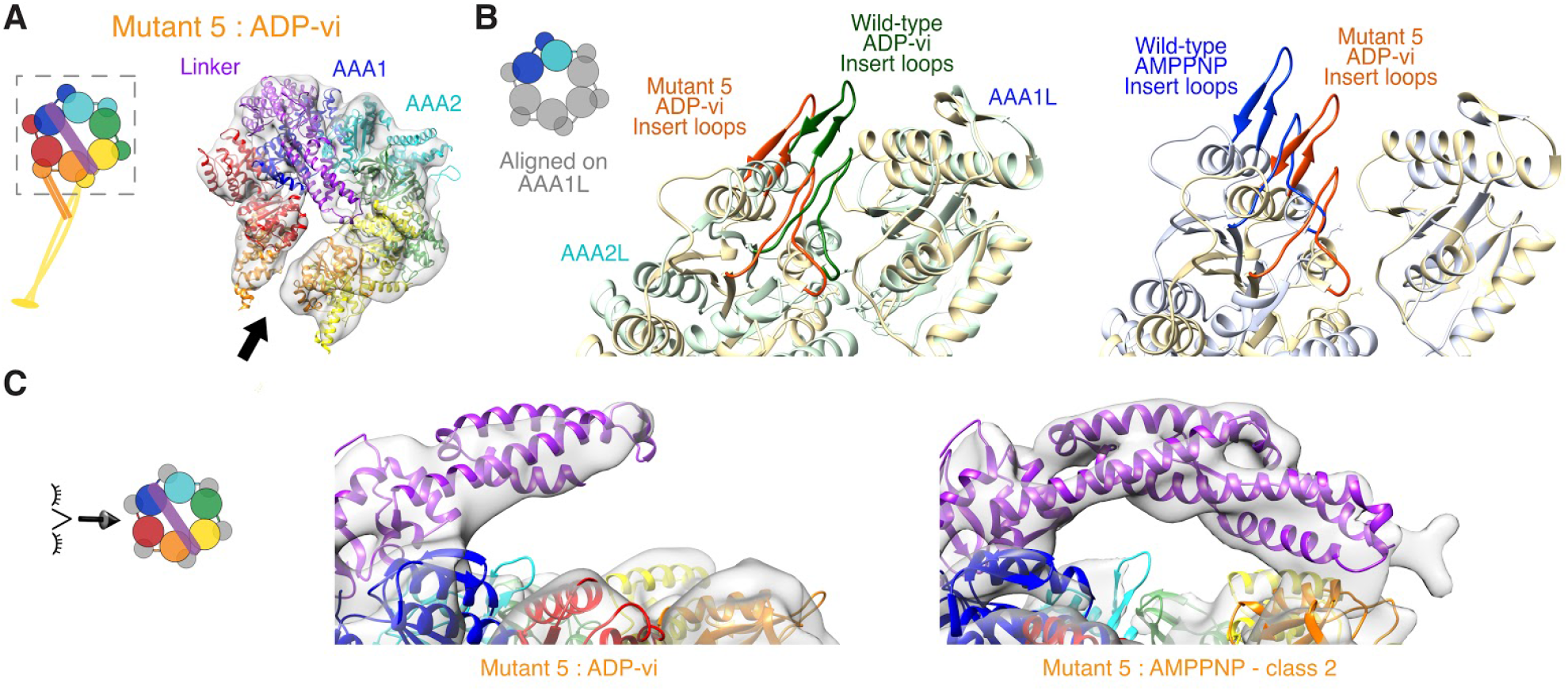
Cryo-EM structure of mutant 5 in the presence of ADP-vanadate shows priming for hydrolysis with unbent linker. (A) Cryo-EM reconstructions and fitted models from 3D classification of the data. The cryo-EM density map is shown as a semi-transparent surface with a fitted model (fit as described in Materials and Methods) as cartoon. Color coding of domains is the same as for Fig. 1. Left: schematic of monomeric dynein construct, box indicates region that was resolved in the cryo-EM map. (B) Close-up view of the AAA1 and AAA 2 interface. The AAA2L inserts (the ‘H2 insert’ and the ‘pre-sensor-I’ (PS-I) insert) are shown in non opaque colors. The structures of human cytoplasmic dynein 2 in the ADP-vi state (green - PDB: 4RH7 (Schmidt et al, 2015)), yeast cytoplasmic dynein in the AMPPNP state (grey - PDB: 4W8F (Bhabha et al, 2014)), and yeast cytoplasmic dynein mutant 5 in ADP-vi (orange - this study) were aligned on AAA1L. Left: box in schematic indicates region of nucleotide pocket. (C) Close-up view of linker of cryo-EM reconstructions and fitted models for mutant 5 in ADP-vi (middle) and in AMPPNP class 2 (right). For the ADP-vi state only the part of the linker with sufficient density was fitted. Left: schematic shows the point-of-view.

To better understand how mutant 5 can be primed for hydrolysis while the linker remains in a straight conformation, we analyzed the AAA domain movements as dynein transitions from the AMPPNP to the ADP-vi state. When both states are aligned on AAA1L, we observe similar domain movements in approximately one-half of the ring surrounding AAA1 (from AAA5s to AAA2L) (Fig. 5A, Supplementary Movie 13), while the domain movements in the other half of the ring, AAA2s to AAA5L, are quite different (Fig. 5B, Supplementary Fig. 9 c). In contrast to the pronounced nucleotide-dependent motions in the AAA2s-AAA5L half of the ring for wild-type dynein, very little motion is observed for these domains in mutant 5 and the mode of movement is different (Fig. 5B). Thus, between the AMPPNP and ADP-vi states, mutant 5 exhibits normal AAA domain movements in one half of the ring (AAA5s-AAA2L), but shows a considerable lack of motion in the other half (AAA2s-AAA5L) (Fig. 5A, B). This result reveals that this stalk mutation uncouples nucleotide-dependent conformational changes in the two halves of the ring. Moreover, these results provide new insight into domain movements of the AAA ring and could explain why mutant 5 shows high ATPase activity but little motility, as will be described in the discussion.

**Figure 5.**
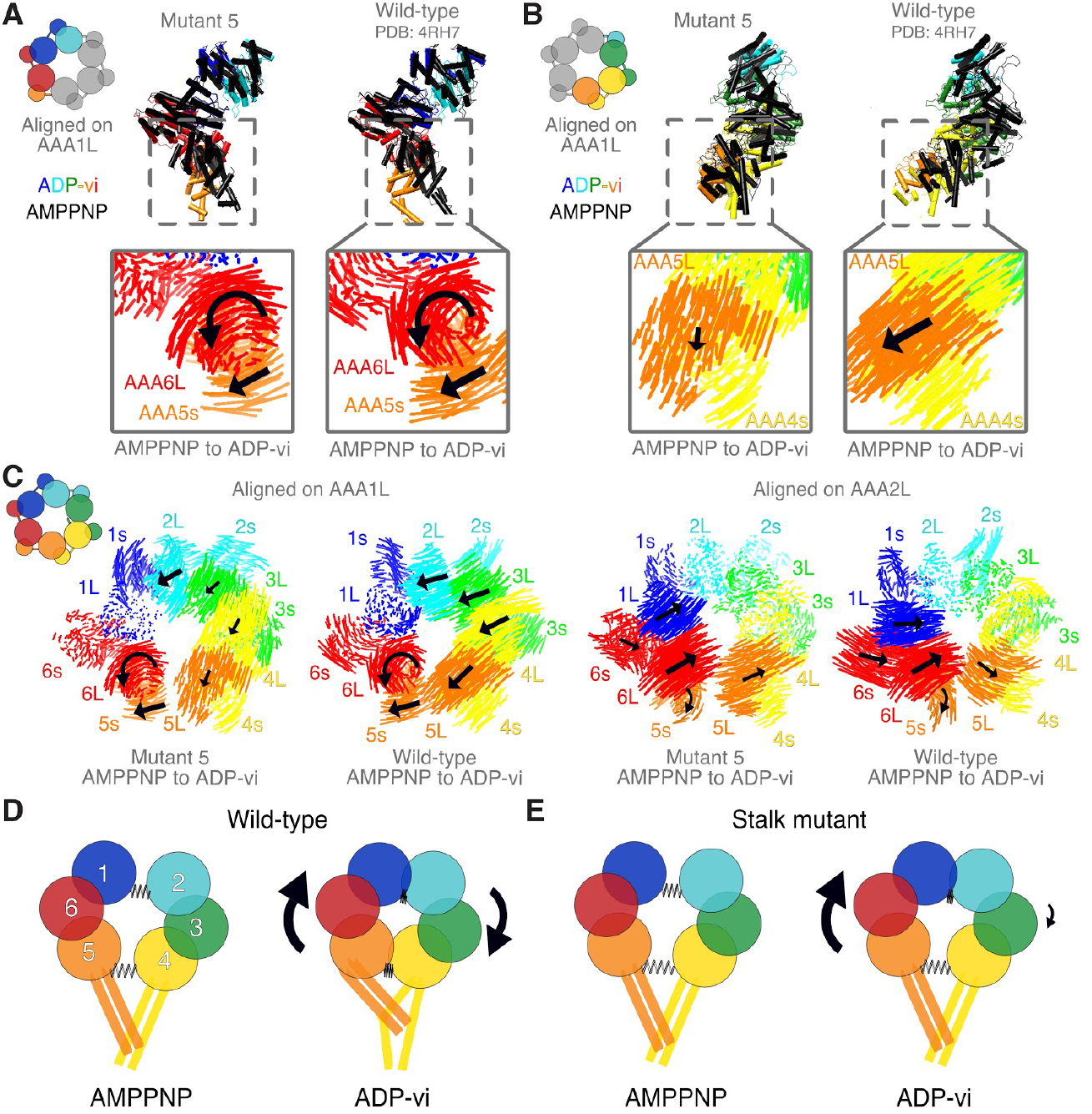
Domain movements in the AAA ring of dynein. (A) Domains AAA5s to AAA2L of wild-type and mutant 5 are shown for the ADP-vi state (color) and the AMPPNP state (black). Box: Visualization of inter alpha carbon distances between AMPPNP and ADP-vi state of mutant 5 and wild-type dynein. Black arrows indicate direction of movement when transitioning from the AMPPNP to the ADP-vi state while the size of the arrow indicates the magnitude of movement. All structures are alignment on AAA1L. (B) Same as in A but for domains AAA2s to AAA5L. (C) Visualization of inter alpha carbon distances between the AAA domains in the AMPPNP and the ADP-vi state of mutant 5 and wild-type dynein for alignments on AAA1L (left) and AAA2L (right). Black arrows indicate direction of movement when transitioning from the AMPPNP to the ADP-vi state while the size of the arrow indicates the magnitude of movement. We removed the linker for clarity. (D) Model for domain movements in the AAA ring of dynein during ATP hydrolysis at AAA1. The AAA ring can be divided into two halves that are connected by two springs. Upon ATP binding / hydrolysis, the gap between AAA1/ AAA2 closes and moves AAA6 / AAA5 which in turn pulls on AAA4 so that the gap between AAA4 / AAA5 closes as well. In addition, this conformational change will pull the buttress and therewith change the stalk registry. (E) In the stalk mutant, the spring between AAA4 / AAA5 does not close upon ATP binding / hydrolysis presumably due to a disruption of the stalk / buttress interface. Moreover, the gap at the AAA4 / AAA5 interface is larger for the stalk mutant in both states, AMPPNP and ADP-vi, than for wild-type in the AMPPNP state. This “loose spring” at AAA4 / AAA5 uncouples these domains from the closure of the AAA1 / AAA2 interface, and this accounts for microtubule independent hydrolysis. (A-C) For wild-type, the structures of human cytoplasmic dynein 2 (ADP-vi state - PDB: 4RH7 (Schmidt et al, 2015)), and the yeast cytoplasmic dynein (AMPPNP state - PDB: 4W8F (Bhabha et al, 2014)) were used. For mutant 5 AMPPNP, we used the class 1 structure.

## Discussion

We have identified mutations in the dynein stalk that show nucleotide-independent weak binding to microtubules and diffusional motion along the microtubule surface. A microtubule-stimulated ATPase assay revealed that these mutants hydrolyze ATP independently of microtubule concentration; two of these mutants are hyperactive and have a basal ATPase activity that is as high as the maximal microtubule-stimulated turnover rate in the wild-type protein. Performing structural analysis on one of these mutants using cryo-electron microscopy, we found that nucleotide-dependent “straight-to-bent” conformational change in the linker domain is inhibited. Moreover, we observed that AAA domain movements in one part of the ring are altered, while the other part of the ring becomes primed for hydrolysis very similarly as in wild-type dynein. These data provide new information on how the microtubule-binding domain (MTBD), stalk, linker, and AAA ring communicate with one another during the ATPase cycle, as discussed below.

### Domain movements in the AAA ring

Dynein is a large and complex allosteric protein that must coordinate the conformations of four independent domains: 1) the AAA ring (consisting of 6 AAA domains), 2) the largely helical linker (which spans over the ring and serves as a mechanical element), 3) the small, globular microtubule-binding domain, and 4) the stalk-buttress apparatus (a pair of antiparallel coiled-coils that extend from the AAA ring and connect via the stalk to the microtubule binding domain). Current structural data suggests that ATP binding to AAA1, with ADP bound at AAA3, drives full AAA ring closure (Schmidt *et al*, 2015; Kon *et al*, 2012; Bhabha *et al*, 2014), which is associated with a large-scale conformational change in the linker and a shift in registry of the two antiparallel coiled coils that affects the affinity of the distal microtubule binding domain. However, the manner in which these different domains communicate with one another is incompletely understood. Previous models for conformational changes in the AAA ring upon ATP binding suggest a rigid body movement of AAA2-AAA4, which propagates as a rotational motion to AAA5s and AAA6L which in turn pull the buttress relative to the stalk (Schmidt *et al*, 2015; Bhabha *et al*, 2014). ATP hydrolysis and/or product release then straightens the linker (thought to be the “power-stroke”), and relaxes the ring back to its original conformation. Another model by Kon et al. (Kon et al. 2012) suggests an important role of the C-terminal domain (C-sequence), located on the surface of the ring opposite to the linker, in connecting AAA1L and AAA5s and triggering a movement of the buttress and, in turn, conformational changes of the stalk and MTBD. Even though the full C-terminal domain is not found in every dynein (Schmidt *et al*, 2015), the H1 alpha helix of the C-sequence that staples the AAA1L/AAA6s and AAA6L/AAA5s blocks together (Kon et al. 2012) appears to be present in virtually all dyneins.

Comparing the conformational states sampled by mutant 5 with the conformational states previously reported for the wild-type protein, we can understand how the stalk mutation in mutant 5 perturbs normal conformational changes in the dynein motor protein, thus provide insight into stalk-mediated conformational changes of the AAA ring.

Our cryo-EM data of mutant 5 revealed wild-type like nucleotide-dependent AAA domain conformational rearrangements in one part of the ring (AAA5s-AAA6-AAA1-AAA2L), but absence /alteration of these conformational changes in the other part of the ring (AAA2s-AAA3-AAA4-AAA5L) (Fig. 5A, B, Supplementary Movie 13). Specifically, movements in AAA5L and AAA4s are much smaller in magnitude, and different in their vector of movement (Fig. 5B, Supplementary Movie 13). This result indicates that the domain movements in the AAA5s-AAA2L block are insulated, at least to some extent, from the rest of the ring and from disruptive mutations in the stalk (Fig. 5C, Supplementary Movie 14). Thus, we hypothesize that the two halves of the ring can undergo two independent modes of conformational change, which require the stalk-buttress apparatus to be properly coupled.

Based on the conformational changes seen in the cryo-EM data for mutant 5, we speculate that ATP binding does not primarily propagate in a clockwise (viewed from the linker side), domino-like manner from AAA1 to AAA6 (Bhabha *et al*, 2014; Schmidt *et al*, 2015; Carter, 2013). Instead, the data for mutant 5 shows bidirectional domain movement around the ATP-bound pocket of AAA1, with a block of AAA5s-AAA6Ls-AAA1L moving towards the AAA1s-AAA2L block upon ATP binding to AAA1 (Supplementary Movie 14). The C-terminal domain might provide the underlying bridging support between AAA5s/AAA6L and AAA6s/AAA1L that allows this block of AAA domains to move in a unified manner, consistent with the proposal of Kon et al. (Kon et al. 2012).

The structural data also emphasize the important role that the stalk-buttress play in coupling the conformational changes in the two halves of the ring. Although we cannot see the precise lesion in the stalk caused by mutant 5 due to flexibility in this region, the downstream effect is an enlarged gap between AAA5s and AAA5L, which we speculate is the underlying cause in the disruption in the allosteric communication within the ring (Fig. 5D, E). Specifically, our data for mutant 5 in the presence of ADP-vanadate suggests that movement of AAA5s/AAA6L towards the nucleotide binding pocket of AAA1 is unable to pull AAA5L/AAA4s with it (and the other half of the ring) (Supplementary Movie 14, Fig. 5C). The interaction between the buttress (emerging from AAA5s) and stalk (emerging from AAA4s) is likely needed for this coordination between the two halves of the ring. Interestingly, the failure to connect AAA5L/AAA4s to the movement of the AAA5s-AAA6Ls-AAA1Ls-AAA2L block severely impacts the nucleotide-dependent movements of the half of the ring from AAA2s-AAA5L. This result suggests that domain movements in this part of the ring are dependent upon the integrity of the stalk-buttress apparatus and its connection to the autonomous nucleotide-driven motions of the AAA5s-AAA6Ls-AAA1Ls-AAA2L block.

This new model suggests why the stalk mutants hydrolyze ATP independently of microtubule concentration. Specifically, the disruption of the stalk-buttress interface allows the buttress and AAA5s/AAA6/AAA1 L to undergo open-closed transitions accompanying ATP binding, hydrolysis and product release, without any regulation by microtubules through the stalk-buttress apparatus (Fig. 5D, E). Interestingly, buttress mutations that presumably disrupt the stalk-buttress interaction (Kon *et al*, 2012) also show high ATPase activity independent of microtubules, similar to our mutants 5 and 13. Thus, we suggest that the stalk-buttress interaction is a key regulator for dynein’s ATPase activity by controlling the coupling between AAA4s/AAA5L and AAA5s/AAA6L and thus the coordination of domain movements in the two halves of the AAA ring.

### Uncoupling of linker bending from robust ATP hydrolysis

Our structural analysis revealed that mutant 5 has uncoupled nucleotide-dependent changes in one half of the AAA ring and ATP hydrolysis from the large conformational change of the linker domain. Previous findings (Bhabha *et al*, 2014; DeWitt *et al*, 2015) suggested that in order for ATP hydrolysis to proceed, the linker must be undocked from the ring, to allow full closure of AAA2L. Thus far, structures in which the AAA ring is primed for hydrolysis (i.e. the gap between AAA2 and AAA1L is closed) have been accompanied by linker bending and docking onto AAA3/2 (Schmidt *et al*, 2015; Bhabha *et al*, 2014). However, our cryo-EM reconstruction of mutant 5 in the presence of ADP-vanadate shows a motor that is primed for hydrolysis with an unbent linker (Fig. 4A, C).

Weak/missing density for the N-terminus of the linker suggests that it is flexible and might be undocked; however, it is clearly not in a bent conformation (Supplementary Fig. 9B). These data therefore suggest that linker bending is not a prerequisite to prime the motor for hydrolysis and that linker bending is necessary for efficient directional motion (since single mutant 5 motors show random bidirectional motion), consistent with other studies (Bhabha *et al*, 2016; Cleary *et al*, 2014). These data further support our model that the hydrolysis cycle arises from autonomous conformational changes within the AAA5s-AAA6Ls-AAA1Ls-AAA2L block and neither require linker bending nor an intact stalk-buttress interface.

Our structural data also allow us to speculate on how linker bending is initiated and why it fails to occur in the mutant 5. In the nucleotide-free and ADP state, the linker forms contacts with AAA5L. Movement of AAA5L upon ATP binding in AAA1 may induce a steric clash with the linker (perhaps with contributions from AAA4L (Schmidt *et al*, 2015)), and linker bending may ensue to minimize such clashes. In mutant 5, AAA5L movement is minimal and thus may be unable to induce the linker steric clash. We therefore speculate that the interface between AAA5s and AAA5L (AAA5s connecting to the AAA1 ATPase site via the AAA5s-AAA6Ls-AAA1Ls-AAA2L block and AAA5L interacting with the linker and AAA4s/stalk/MTBD) may be a critical region for coordinating ATPase activity (AAA1), microtubule binding (stalk/MTBD), and mechanics (linker) in the dynein motor domain (Supplementary Movie 14).

## Materials and Methods

### Bioinformatic analysis of dynein sequences

The detailed process is described in Supplementary Note 1. Briefly, we used 677 unique axonemal and cytoplasmic dynein heavy chain sequences from 229 fully sequenced eukaryotic genomes, which we received from Christian Zmasek, <u>Godzik lab</u>, Burnham. This data set was pruned based on well defined criteria as listed in Supplementary Note 1 and analyzed using Jalview (Waterhouse *et al*, 2009). Remaining sequences were aligned using MAFFT (Katoh *et al*, 2002) in the Bioinformatic Toolkit (Alva *et al*, 2016) and mutations in the stalk were manually identified by comparing sequences in Jalview. All alignment files are available as Supplementary Material.

### Yeast strains used in this study

Recombinant *S.cerevisiae* cytoplasmic dynein (Dyn1) truncated at the N-terminus (1219-4093 aa) was used in this study. All constructs used in this study are listed in Supplementary Table 1. Dimeric constructs are based on VY208 and were created by artificially dimerization through an N-terminal GST-tag (Reck-Peterson *et al*, 2006) and tagged with a HaloTag (Promega) at the C-terminus as well as a GFP at the very N-terminus. Monomeric constructs (VY137) are GFP tagged at the N-terminus. Stalk mutations were inserted by homologous recombination as previously described (Reck-Peterson *et al*, 2006).

### Protein expression and purification

Dynein was expressed and purified as previously described (Reck-Peterson *et al*, 2006). Monomeric and dimeric constructs were further purified by gel filtration on a GE Healthcare Superdex 200 10/300GL and a GE Healthcare Superose 6 10/300GL column, respectively in dynein gel filtration buffer (50 mM K-Ac, 20 mM Tris, pH 8.0, 2 mM Mg(Ac)_2_, 1 mM EGTA, 1 mM TCEP, and 10% glycerol) and flash frozen afterwards. The ‘cysteine-light’ human ubiquitous kinesin-1 dimer E215C K490 construct was cloned and purified as previously described (Mori *et al*, 2007; Tomishige *et al*, 2006b). Following dialysis the E215C K490 construct was reacted for 4 h at 4°C with Cy3-maleimide (GE Healthcare, PA13131) at a motor/Cy3 dye ratio of 1:10 as previously described (Tomishige *et al*, 2006b). The unreacted maleimide dyes were then quenched with 1 mM dithiothreitol (DTT). Afterwards the kinesin was purified by gel filtration over a S200 10/300GL column (GE Healthcare) in kinesin gel filtration buffer (25 mM Pipes (pH 6.8), 2 mM MgCl_2_, 200 mM NaCl, 1 mM EGTA, 1 mM DTT, and 10% sucrose) and then flash frozen.

### Microtubule preparation

Tubulin was purified and polymerized as previously described (McKenney *et al*, 2014). For single-molecule motility assays unlabeled tubulin, biotinylated tubulin, and fluorescent tubulin were mixed at an approximate ratio of 20:2:1 in BRB80 (80 mM Pipes (pH 6.8), 1 mM EGTA, and 1 mM MgCl_2_). For the gliding assay unlabeled tubulin and fluorescent tubulin were mixed at an approximate ratio of 20:1 in BRB80. For tubulin that was used in the ATPase assay as well as the microtubule affinity assay only unlabeled tubulin was used. We added 1 mM GTP to all polymerization reactions. Then the mixtures were incubated for 15 min in a 37°C water bath. 20 *μ*M of Taxol (Sigma, T1912) was added afterwards and the mixture was incubated for 2 more hours at 37°C. Before usage, microtubules were spun over a 25% sucrose cushion in BRB80 at ~160,000 g for 10 min in a tabletop centrifuge.

### Gliding and single-molecule motility assay

We made custom flow chambers using laser-cut double-sided adhesive sheets (Soles2dance, 9474-08×12 - 3M 9474LE 300LSE). We used glass slides (Thermo Fisher Scientific, 12-550-123), and coverslips (Zeiss, 474030-9000-000). We cleaned the coverslips in a 5% v/v solution of Hellmanex III (Sigma, Z805939-1EA) at 50°C overnight and then washed them extensively with Milli-Q water. The flow-cells were assembled in a way that each chamber holds approximately 10 *μ*l.

Every data collection was carried out at room temperature (~23°C) using a total internal reflection fluorescence (TIRF) inverted microscope (Nikon Eclipse Ti microscope) equipped with a 100× (1.45 NA) oil objective (Nikon, Plan Apo Λ). We used an Andor iXon 512×512 pixel EM camera, DU-897E and a pixel size of 159 nm. Dynein (always as dimer and either labeled with GFP only or with GFP and a Halo488 dye (Promega, G1001)) was excited with a 488 nm laser (Coherent Sapphire 488 LP, 150 mW), kinesin with a 561 nm laser (Coherent Sapphire 561 LP, 150 mW), and microtubules with a 640 nm laser (Coherent CUBE 640-100C, 100 mW). For the gliding assay, images were recorded with 100 ms exposure time and a 2 sec frame rate for MTs and a 100 msec frame rate for kinesin. For the single-molecule assay of dynein, we used 100 msec exposures and a 2 sec frame rate and a 100 msec frame rate for kinesin. The acquisition software was *μ*Manager (Edelstein *et al*, 2010) 2.0 and data was analyzed in ImageJ (Schneider *et al*, 2012).

For the gliding assay, we first added 10 μl of GFP antibody (Abcam, ab1218) and incubated for 5 min. Then we washed with 20 μl of DAB with 2 mg/ml *β*-casein and 0.4 mg/ml *κ*-casein. We then added 10 μl of dynein and incubated for another 5 min which was followed by an additional wash with 20 μl of DAB with 2 mg/ml *β*-casein and 0.4 mg/ml *κ*-casein. Next, we added 10 μl of polymerized microtubules and incubated for 5 min. Then we washed with 30 μl of DAB with 2 mg/ml *β*-casein and 0.4 mg/ml *κ*-casein. Finally, 10 μl of DAB with kinesin, 0.4 mg/ml *κ*-casein, 10 μM Taxol, 1 mM Mg-ATP, and the PCA/PCD/Trolox oxygen scavenging system (Aitken *et al*, 2008) was added.

Prior to the single-molecule motility assays dynein was labeled with Halo488 dye (Promega, G1001) as previously described (Bhabha *et al*, 2014). Briefly, dynein constructs were mixed with 20 μM Halo Alexa488 dye and incubated on ice for 10 min and a PD MiniTrap G-25 column (GE Healthcare) equilibrated with dynein gel filtration buffer was used to remove excess dye afterwards.

The flow chambers for the single-molecule motility assay were prepared as previously described (Yildiz & Vale, 2015). Briefly, we first added 10 μl of 5 mg/ml Biotin-BSA in BRB80 and incubated for 2 min. Then we washed with 20 μl of BRB80 with 2 mg/ml *β*-casein (Sigma, C6905), 0.4 mg/ml *κ*-casein (Sigma, C0406). Afterwards we added 10 μl of 0.5 mg/ml Streptavidin in PBS for a 2 min incubation. Next, we again washed with 20 μl of BRB80 with 2 mg/ml *β*-casein, and 0.4 mg/ml *κ*-casein. This was followed by the addition of 10 μl of polymerized microtubules and a 5 min incubation. Then we washed with 30 μl of DAB (50 mM K-Ac, 30 mM HEPES, pH 7.4, 2 mM Mg(Ac)_2_, 1 mM EGTA) with 2 mg/ml *β*-casein, 0.4 mg/ml *κ*-casein, and 10 μM Taxol. Finally we added 10 μl of dynein and kinesin in DAB with 0.4 mg/ml *κ*-casein, 10 μM Taxol, 1 mM Mg-ATP, and the PCA/PCD/Trolox oxygen scavenging system (Aitken *et al*, 2008). In the single-molecule assay where ATP was omitted, the final solution contained 10 μl of dynein in DAB with 0.4 mg/ml *κ*-casein, 10 μM Taxol, and the PCA/PCD/Trolox oxygen scavenging system (Aitken *et al*, 2008).

### ATPase assay

The ATPase assays were carried out in DAB (50 mM K-Ac, 30 mM HEPES, pH 7.4, 2 mM Mg(Ac)2, 1 mM EGTA) as follows. We mixed dynein (monomeric for all constructs) to a final concentration of 10-20 nM with 2 mM Mg-ATP (Sigma), 0.2 mM NADH (Sigma), 1 mM phosphoenolpyruvate (Sigma), 0.01 U pyruvate kinase (Sigma), 0.03 U lactate dehydrogenase (Sigma), 10 μM Taxol, 1mM DTT, and 0-5 μM microtubules in DAB. Absorbance at 340 nm was continuously measured in an Eppendorf Spectrophotometer (UV-Vis BioSpectrometer) and the data was fit to the following equation (Bhabha *et al*, 2014) using an excel curve fitting routine: 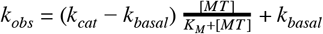.

### Microtubule affinity assay

The microtubule affinity assays were carried out in DAB (50 mM K-Ac, 30 mM HEPES, pH 7.4, 2 mM Mg(Ac)_2_, 1 mM EGTA) as follows. We mixed dynein (monomeric for all constructs) to a final concentration of approx. 50 nM with 10 μM Taxol, 1 mM DTT, and 0-8 μM microtubules in DAB. For the measurements with ATP we added 5 mM Mg-ATP (Sigma) and for the experiment with AMPPNP we added 5 mM Mg-AMPPNP (Sigma). After a 3 min incubation at room temperature the samples were spun over a 25% sucrose cushion in DAB at ~160,000 *g* for 10 min in a tabletop centrifuge. The concentration of dynein in the supernatant (unbound) and in the pellet (bound) was determined by measuring the intensity of the N-terminal GFP on a Typhoon (GE Healthcare) and the data was fit to the following equation using an excel curve fitting routine: 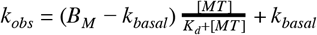 (B_M_ maximum binding, K_d_ dissociation constant).

### Electron microscopy data collection

For negative stain, data for mutants 5 (monomer) was collected on a Tecnai F20 microscope with a Tietz F416 CMOS detector at the New York Structural Biology Center (NYSBC). Leginon software (Suloway *et al*, 2005) was used for the semi-automated collection of 825 images at a magnification of x62,000 and a pixel size of 3 Å per pixel. For cryo-EM data collection, 1200 movies of mutant 5 (monomer) mixed with 2 mM AMPPNP were recorded with SerialEM (Mastronarde, 2005) at 300 kV on a Titan Krios (FEI) equipped with a K2 summit camera (Gatan) at 0.655 Å per pixel in super-resolution mode at Janelia Research Campus. Another 664 movies of the same mutant (mutant 5 - monomer) mixed with 2 mM ATP and 2 mM vanadate were recorded with SerialEM at 200 kV on a Arctica (FEI) equipped with a K2 summit camera (Gatan) at 0.578 Å per pixel in super-resolution mode at New York University.

### Electron microscopy data processing and analysis

For the images of the negatively stained sample, particles were selected using DoG picker (Voss *et al*, 2009) in APPION (Lander *et al*, 2009), then extracted in Relion 2.1.0 (Scheres, 2012) into boxes of 180×180 pixels, leading to 156,199 boxes for mutant 5. A round of 2D classification was performed to remove junk and noisy particles, leading to 54,913 particles selected. Subsequent image processing steps were carried out using CryoSPARC (Punjani *et al*, 2017). After having generated an ab-initio model, those particles were used to generate eight 3D classes. Because of the similarity between all those classes, a final round of 3D refinement was completed using all of the particles.

For the cryo-EM images (see also Supplementary Table 5), the movies of mutant 5 with 2 mM AMPPNP were first aligned and binned to 1.31 Å per pixel with MotionCor2 v1.0.5 (Zheng *et al*, 2017), and then the contrast transfer function parameters were estimated with GTCF 1.06 (Zhang, 2016). The particles were picked automatically in Relion 2.1.0 (Scheres, 2012) using a Gaussian blob as a reference and further processing was done in CryoSPARC (Punjani *et al*, 2017). Out of the 310,085 regions automatically picked, 136,056 were kept after evaluation of 2D classes. Two ab-initio models were first generated in CryoSPARC, and the best one was used in a 4-class 3D heterogeneous refinement.

Then, two 3D homogeneous refinements were completed: one with class 3 (here refered to as class 2 - with 29% of remaining particles and with a resolution of 7.6 Å), and another one (here refered to as class 1 - with 71% of remaining particles and with a resolution of 7.7 Å) with the three other classes which looked very similar and were therefore combined before refinement. The final maps were then filtered for display using a B-factor of −400. For modelling, we used PDB 4W8F as a reference. The PDB file was split into 13 domains (small and large subdomains for each AAA domain, and the linker) and, for each of those domains, we simultaneously fit all 13 subdomains into the map using UCSF Chimera (Pettersen *et al*, 2004). We noticed that the rigid body of the buttress region in class 2 map did not perfectly fit the densities (Supplementary Fig. 7 c). This model was therefore subjected to the real_space_refine algorithm in PHENIX (Adams *et al*, 2010) using 2 cycles and 100 iterations to optimize the fit. Figures and movies were generated with the UCSF Chimera package or the Pymol Molecular Graphics System (version 2.0, Schröodinger, LLC). For the images of mutant 5 with 2 mM ATP and 2 mM vanadate acquired on the Arctica, a similar process was followed. First aligned and binned to 1.31 Å per pixel with MotionCor2 v1.0.5 (Zheng *et al*, 2017), and the contrast transfer function parameters estimated with GCTF 1.06 (Zhang, 2016). A first round of auto-picking was conducted in Relion 2.1.0 (Scheres, 2012) using a Gaussian blob as a reference. Two of the resulting classes were then used as template for a round of reference-based auto-picking. Further processing was also conducted in CryoSPARC (Punjani *et al*, 2017). Out of 35,565 picked particles, 32,442 particles were kept for the generation of ab-initio models and a 4 and a 5-class heterogeneous refinement were tried. One class with 8,653 particles lead to a clear dynein 3D model that we refined to 9.2 Å and finally filtered for display using a B-factor of −400. Another class at 17 Å seemed to show only the AAA domains while the linker could not be seen. The model of the 9.2 Å model was constructed as for mutant 5 in apo-state, using rigid body docking of domains from PDB 4W8F, but was not further refined in PHENIX due to its lower resolution.

### Figure preparation

Figures and graphs were created using Pymol (version 2.0 Schröodinger, LLC) and Chimera (Pettersen *et al*, 2004) (structure representation, ImageJ (Schneider *et al*, 2012) (light microscopy data), Jalview (Waterhouse *et al*, 2009) (sequence analysis and representation), Affinity designer (version 1.6.1, Serif (Europe) Ltd) and Python (version 2.7, Python Software Foundation).

### Statistics

For each result obtained, the inherent uncertainty due to random or systematic errors and their validation are discussed in the relevant sections of the manuscript. Details about the sample size, number of independent calculations, and the determination of error bars in plots are included in the figures and figure captions.

### Data availability

Density maps for the structures were deposited in the Electron Microscopy Data Bank under accession codes EMD-7829 (Mutant 5 in the presence of AMPPNP, Class 1), EMD-7830 (Mutant 5 in the presence of AMPPNP, Class 2), and EMD-9386 (Mutant 5 in the presence of ADP-vanadate). The sequence alignment files used to create the mutants and all other data are available from the corresponding author upon request.

## Supporting information

Supplementary_Datasets_Sequence-alignments

## ACKNOWLEDGMENTS

Cryo EM data were collected on the Titan Krios (“Krios 2”) at Janelia Research campus, and Talos Arctica at the NYU cryo EM core facility. For cryo-EM on the Krios, we thank Hui-Ting Chou and Zhiheng Yu at the HHMI Janelia Research Campus for assistance in microscope operation and data collection. For cryo-EM on the Arctica, we thank Zheng Liu for assistance and data collection. We are grateful to J. Sheu-Gruttadauria and Iris Grossman-Haham for critical discussions of the manuscript. We would like to thank Christian Zmasek at the Burnham Institute for the initial sequence alignment file. We thank Nico Stuurman and Walter Huynh for their assistance and advice in microscopy. Some of our work was performed at the Simons Electron Microscopy Center and National Resource for Automated Molecular Microscopy located at the New York Structural Biology Center. We thank Kelsey Jordan at the New York Structural Biology Center for assistance with data collection of negatively stained samples. For EM data processing, this work has utilized computing resources at the High-Performance Computing Facility at NYU Langone Medical Center. We thank Martin Ossowski and his HPC team as well as Joe Katsnelson for EM data processing support. The authors gratefully acknowledge funding support from the NIH National Institute of General Medical Sciences: R00GM112982 (G.B.), R01GM097312 (R.D.V.), Damon Runyon Cancer Research Foundation DFS-20-16 (G.B.), Howard Hughes Medical Institute (R.D.V.) and the UCSF Discovery Fellowship (S.N.). Some of this work was performed at the Simons Electron Microscopy Center and National Resource for Automated Molecular Microscopy located at the New York Structural Biology Center, supported by grants from the Simons Foundation (349247), NYSTAR, and the NIH National Institute of General Medical Sciences (GM103310).

## AUTHOR CONTRIBUTIONS

G.B., S.N. and R.D.V. conceived the research. G.B., S.N., N.C. and N.Z. performed experiments and collected data. G.B., S.N., N.C. and R.D.V. analyzed data. G.B., S.N., and R.D.V. wrote the manuscript. All authors edited the manuscript.

## CONFLICT OF INTEREST

The authors declare that they have no conflict of interest.

