## Supplementary_Information for "Coupling of ATPase activity, microtubule binding and mechanics in the dynein motor domain"

*for*

<u>This document includes:</u>

Supplementary Information Figures (Supplementary Fig. 1 - 9)

Supplementary Information Notes (1-3)

Supplementary Information Tables (Supplementary Table 1 - 5)

Supplementary Information Movies (Supplementary Movie 1 - 14)

References for the Supplementary Information

### Supplementary Figures

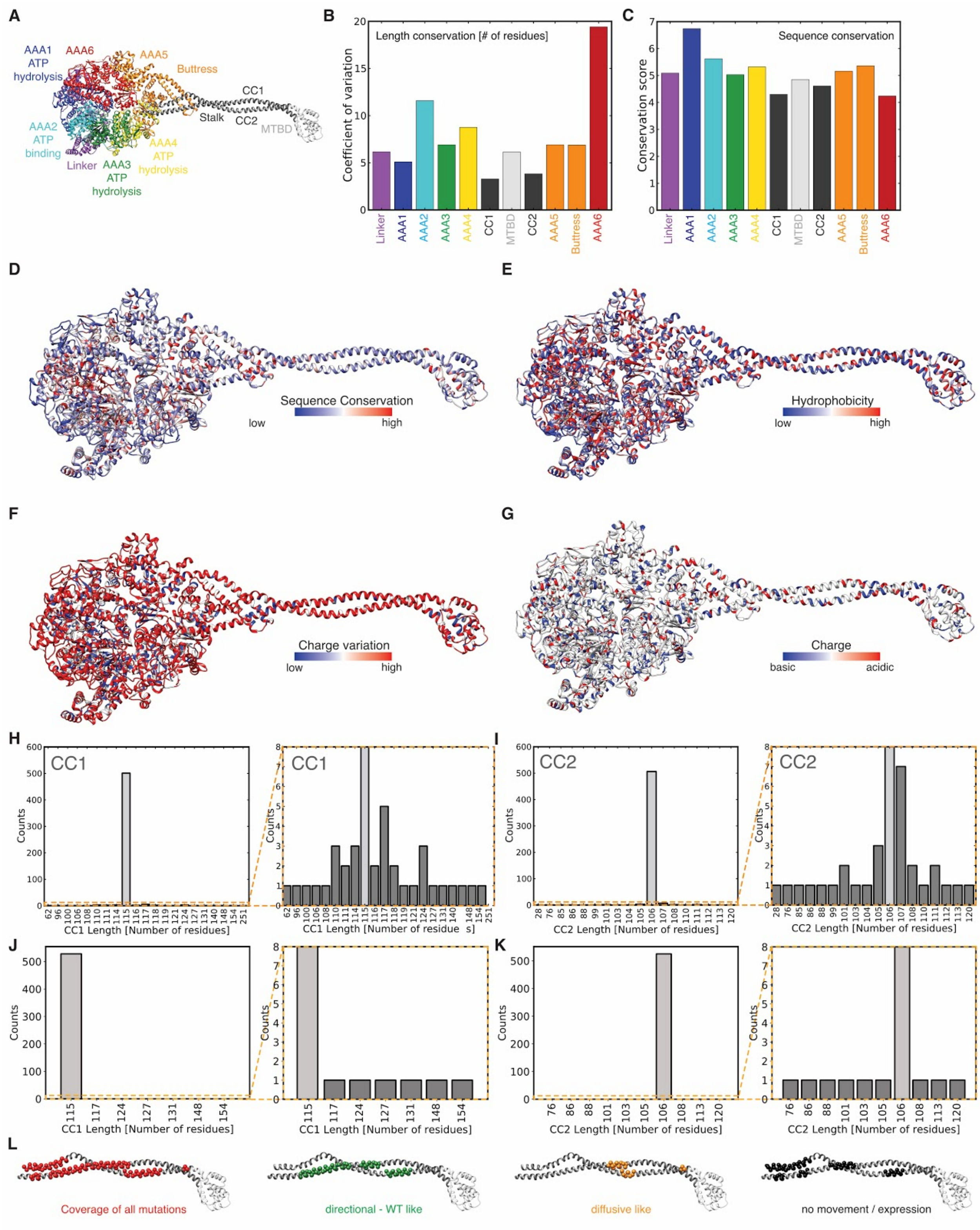

**Supplementary Figure 1 |** Sequence analysis and conservation in the dynein motor domain. (A) Structure and domain organization of the motor domain of cytoplasmic dynein (PDB: 4rh7 (Schmidt *et al*, 2015)). (B) Length (number of residues) conservation of domains of the dynein motor domain. (C) Sequence conservation of domains of the dynein motor domain. Conservation score from Jalview (Livingstone & Barton, 1993; Waterhouse *et al*, 2009) is shown for each domain. The conservations in B and C are based on 534 different sequences that were curated as described in Supplementary Note 1. (D) Sequence conservation, (E) Conservation of hydrophobic residues, (F) Charge variation (how many residues at the same position among different sequences switch between D/E and H/K/R), and (G) Conservation of charge where basic residues (D/E) are in blue and acidic residues (H/K/R) are in red. The conservations shown in D-G are based on 534 different sequences that were curated as described in Supplementary Note 1. (H, I) Histogram showing the length distribution of (H) CC1 and (I) CC2 of the dynein stalk among 534 sequences with initial sequence data (used to derive mutants) as described in Supplementary Note 1. Orange box indicates area that is magnified on the right. (J) Histogram showing the length distribution of CC1 of the dynein stalk among 534 sequences that were updated based on most recent sequencing reads in various data bases as described in Supplementary Note 1. Orange box indicates area that is magnified on the right, showing a handful of outlier sequences with different stalk lengths. (K) Same as in J but for CC2. (L) Coverage of all mutations (red) generated in the yeast dynein background based on our sequence analysis (most left) mapped onto the structure of human cytoplasmic dynein 2 stalk (PDB: 4rh7(Schmidt *et al*, 2015)). Regions of individual mutants are shown in Supplementary Fig. 3. Positions of insertions/deletions that showed ‘Directional - WT like’ (green), ‘Diffusive like’ (orange) movement, and ‘No movement / No expression’ (black) mapped onto the stalk (classification as shown in Fig. 1).

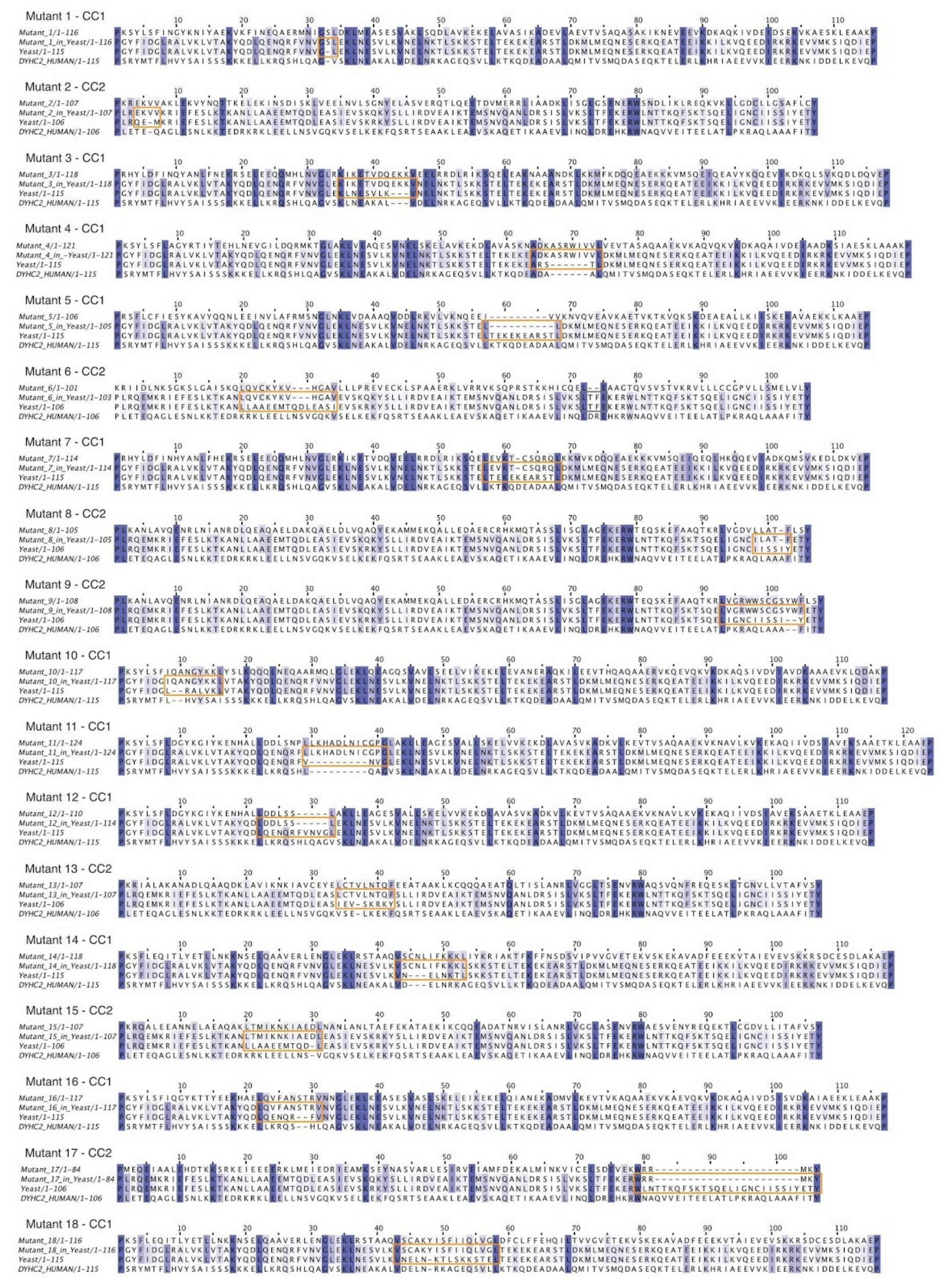
Mutant 1 - CC1
Mutant 2 - CC2
Mutant 3 - CC1
Mutant 4 - CC1
Mutant 5 - CC1
Mutant 6 - CC2
Mutant 7 - CC1
Mutant 8 - CC2
Mutant 9 - CC2
Mutant 10 - CC1
Mutant 11 - CC1
Mutant 12 - CC1
Mutant 13 - CC2
Mutant 14 - CC1
Mutant 15 - CC2
Mutant 16 - CC1
Mutant 17 - CC2
Mutant 18 - CC1

**Supplementary Figure 2 |** Sequence alignments of the panel of stalk mutants. For each of the 18 mutants we compare the sequence of the species with insertion or deletion (top sequence), mutant created in yeast dynein background (second from the top), yeast dynein wild-type (second from bottom), and human cytoplasmic dynein 2 (bottom). Orange boxes highlight area of mutation. Note: Grey box in sequence alignment for mutant 6 shows second position of mutation for mutant 6 which was not created. Sequence conservation is indicated from white (not well conserved) to blue (highly conserved). For more details on how sequences were aligned and how mutants were selected see Supplementary Note 1.

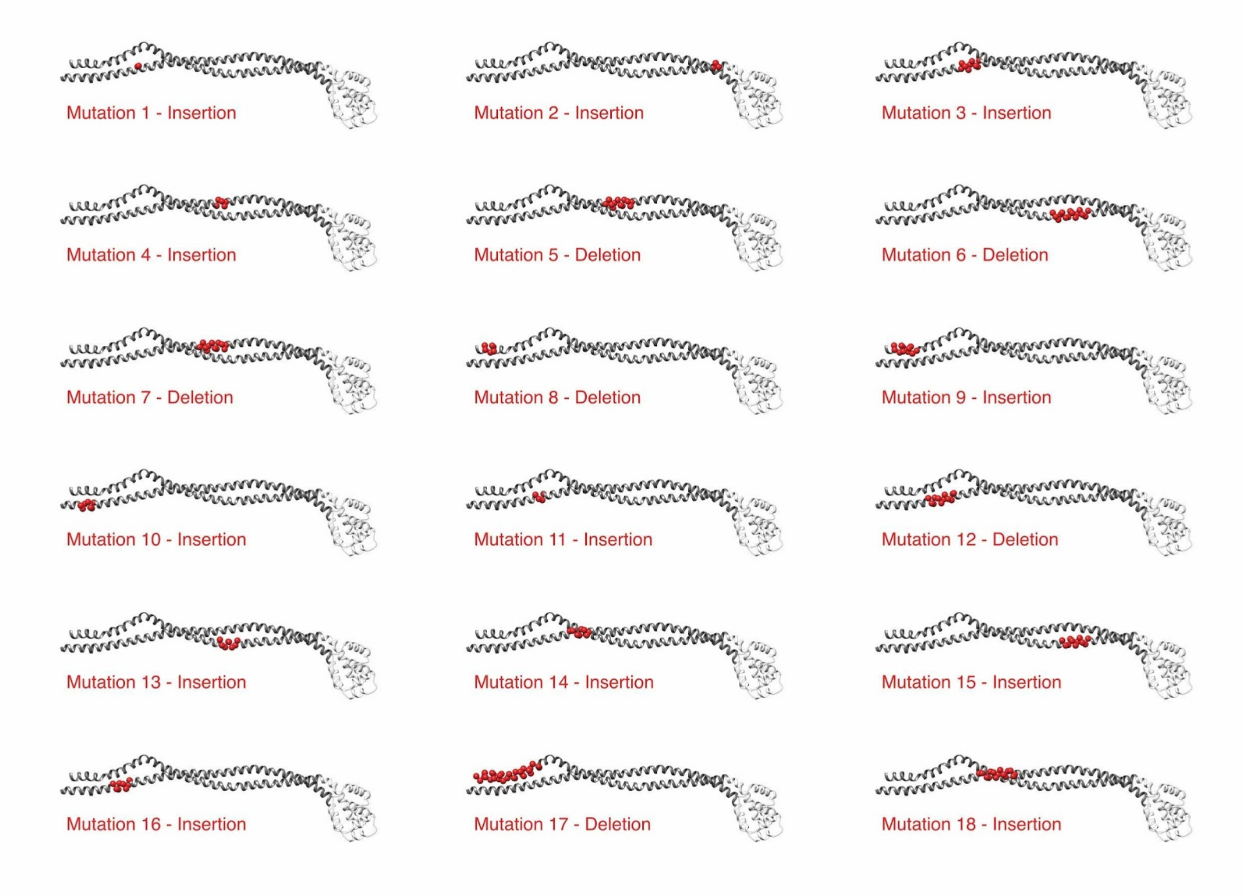


**Supplementary Figure 3 |** Position of all 18 insertion or deletion mutants mapped onto the structure of human cytoplasmic dynein 2 stalk (PDB: 4rh7 (Schmidt *et al*, 2015)). Red spheres show residues that were altered in the stalk to either create an insertion or deletion.

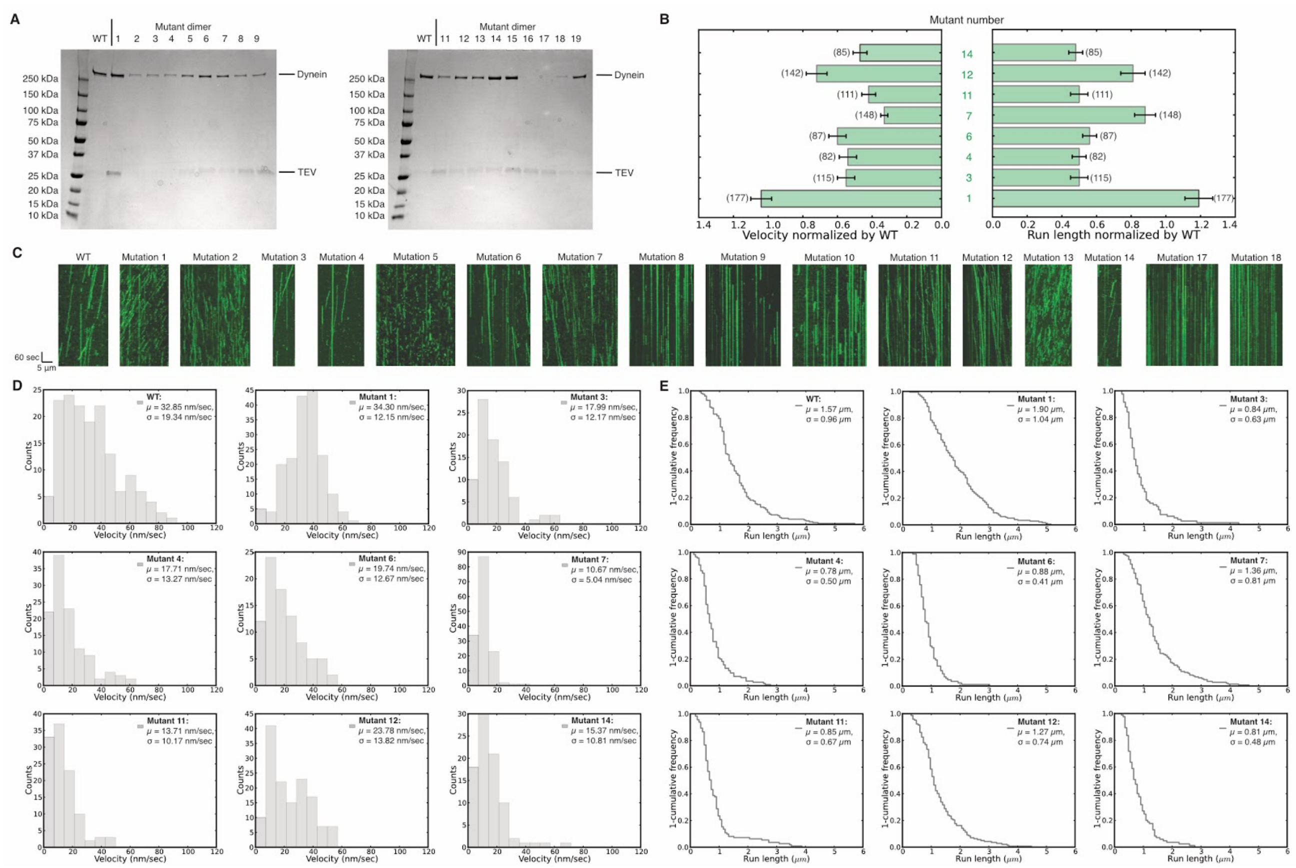


**Supplementary Figure 4 |** Example kymographs and single-molecule motility properties of stalk mutants. (A) Purified dynein, wild-type and mutants, after affinity purification shown by PAGE. No dynein band is visible for mutant 15 and 16 indicating that they did not express. For some constructs residual TEV, which was used the cleave the dynein of beads during the affinity purification (see Materials and Methods), is visible. All constructs that were used for assays other than the single-molecule motility assay were further purified by size exclusion chromatography, which removed the residual TEV entirely (see Materials and Methods). (B) Velocity and run length of 'Directional - WT like' motors normalized by wild-type dynein. Error bars show standard deviation and number in brackets indicates the number of motors quantified. Data used for quantification shown in D and E. (C) Example kymographs showing different types of movement as classified in Fig. 1. Kymographs for mutant 15 and 16 are not shown since they did not express. (D) Velocity histogram with average velocity ($\mu$) and its standard deviation ($\sigma$) for wild-type and 'Directional - WT like' mutants. (E) A '1-cumulative frequency distribution plot' of run length with average length ($\mu$) and its standard deviation ($\sigma$) for wild-type and 'Directional - WT like' mutants. Quantification is based on three repeats of a single preparation.

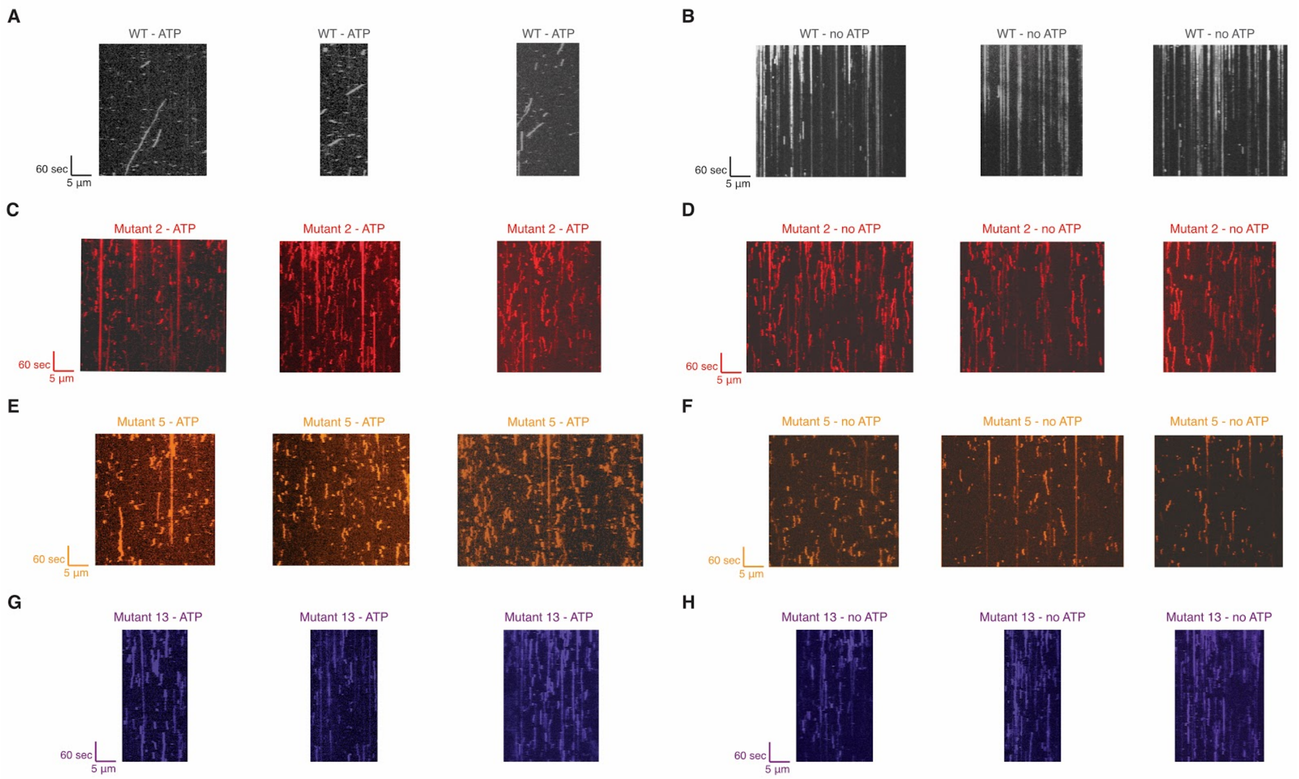


**Supplementary Figure 5 |** Kymographs of wild-type and diffusive mutants with and without ATP. (A, C, E, G) Kymographs of wild-type (grey), mutant 2 (red), mutant 5 (orange), and mutant 13 (purple) with ATP. (B, D, F, H) Kymographs of wild-type (grey), mutant 2 (red), mutant 5 (orange), and mutant 13 (purple) without ATP.

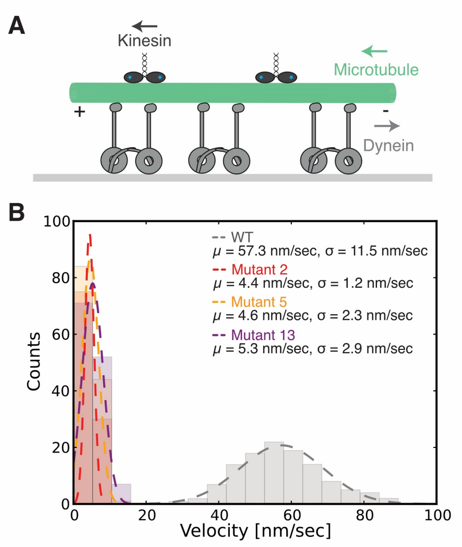


**Supplementary Figure 6 |** Gliding assay shows slow directional movement for mutants 2, 5, and 13. (A) Schematic of modified gliding assay. Dyneins (dark grey) are immobilized on microscope slide (light grey) and can translocate microtubule (green). Plus end directed kinesins (dark blue) move on top of microtubule to mark directionality. (B) Histogram of gliding velocities of wild-type (grey, n=116), mutant 2 (red, n=105), mutant 5 (orange, n=129), and mutant 13 (purple, n=130) with average velocity ($\mu$) and its standard deviation ($\sigma$). Example movies of microtubule gliding for all four constructs are shown in Supplementary Movies 9-12. Data of one dynein preparation is shown but a total of three repetitions of different dynein preparations resulted in very similar velocities.

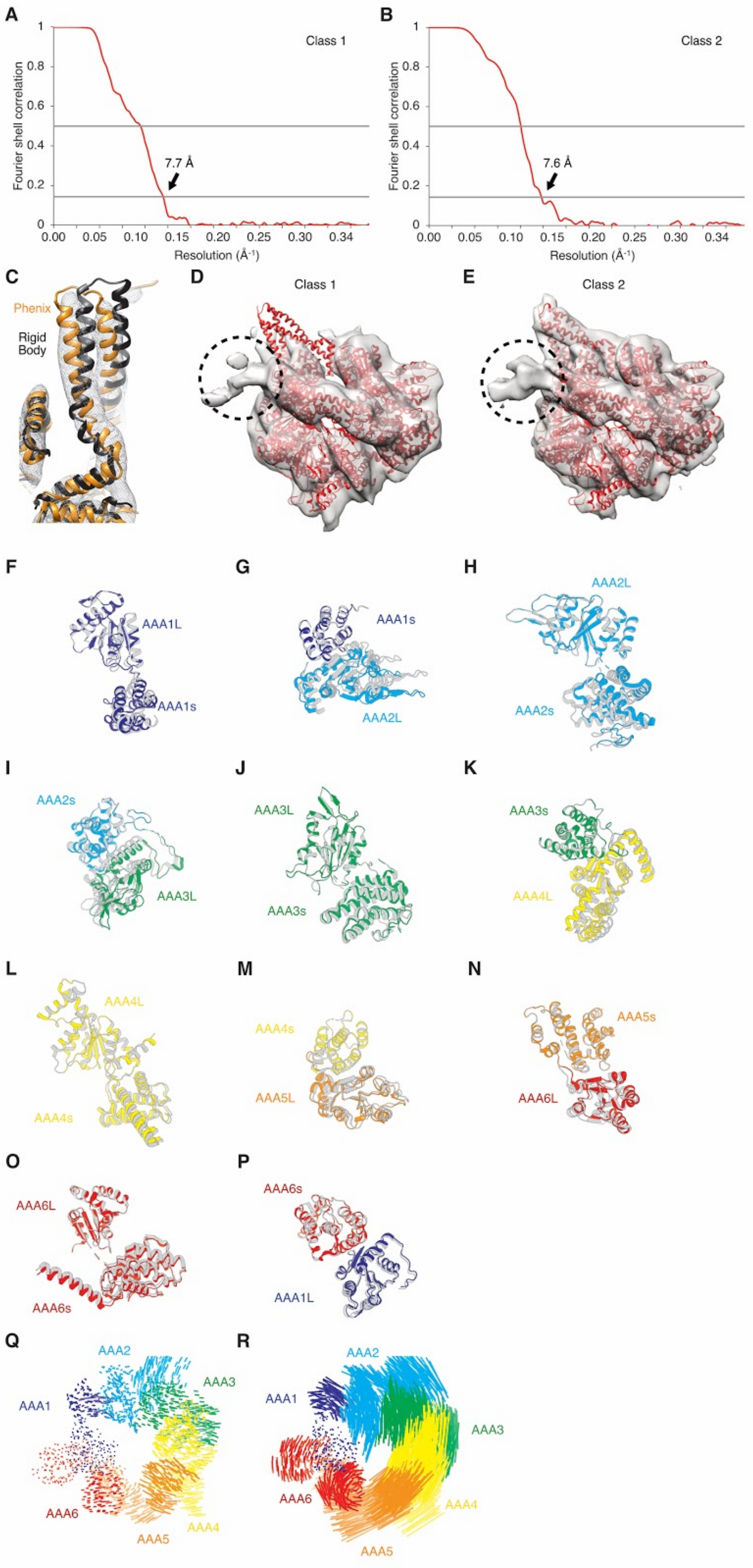
A
Class 1
Fourier shell correlation
7.7 Å
Resolution (Å⁻¹)
B
Class 2
Fourier shell correlation
7.6 Å
Resolution (Å⁻¹)
C
Phenix
Rigid Body
D
Class 1
E
Class 2
F
AAA1L
AAA1s
G
AAA1s
AAA2L
H
AAA2L
AAA2s
I
AAA2s
AAA3L
J
AAA3L
AAA3s
K
AAA3s
AAA4L
L
AAA4L
AAA4s
M
AAA4s
AAA5L
N
AAA5s
AAA6L
O
AAA6L
AAA6s
P
AAA6s
AAA1L
Q
AAA2
AAA3
AAA1
AAA6
AAA5
AAA4
R
AAA2
AAA1
AAA3
AAA6
AAA4
AAA5

**Supplementary Figure 7 |** Cryo-EM analysis for class 1 and class 2 of mutant 5 in the presence of AMPPNP. Plot of Fourier Shell Correlation (FSC) for (A) class 1 and (B) class 2. (C) Cryo-EM density for the buttress region of class 2 with rigid body and flexibly fit models. The rigid body fit of AAA5L into the density clearly showed that some rearrangement of the buttress had occurred. Flexible fitting in Phenix resulted in a model that fit the density in the buttress region significantly better. (D, E) Cryo-EM reconstruction of class 1 and 2 showing unfiltered maps with AMPPNP-bound crystal structure (PDB: 4W8F, red) shown for reference. Black dotted-circle indicates position of a GFP tag at the N-terminus of the linker, which is better defined in class 2. (F-P) Domain movements between class 1 and class 2 of mutant 5 in AMPPNP cryo-EM data. In every panel the top domain is aligned, showing movement between that domain and the next. Class 1 is colored and class 2 is grey. (F) Movement between AAA1L and AAA1s. (G) Movement between AAA1s and AAA2L. (H) Movement between AAA2L and AAA2s. (I) Movement between AAA2s and AAA3L. (J) Movement between AAA3L and AAA3s. (K) Movement between AAA3s and AAA4L. (L) Movement between AAA4L and AAA4s. (M) Movement between AAA4s and AAA5L. (N) Movement between AAA5s and AAA6L. (O) Movement between AAA6L and AAA6s. (P) Movement between AAA6s and AAA1L. (Q) Visualization of inter alpha carbon distances between class 1 and class 2 of mutant 5 in the AMPPNP state after alignment on AAA1L as seen from the top. We removed the linker for clarity. (R) Visualization of inter alpha carbon distances between class 1 of mutant 5 in the AMPPNP state and the cryo-EM model of yeast dynein in the presence of ADP-vi (Bhabha *et al*, 2014). We removed the linker for clarity.

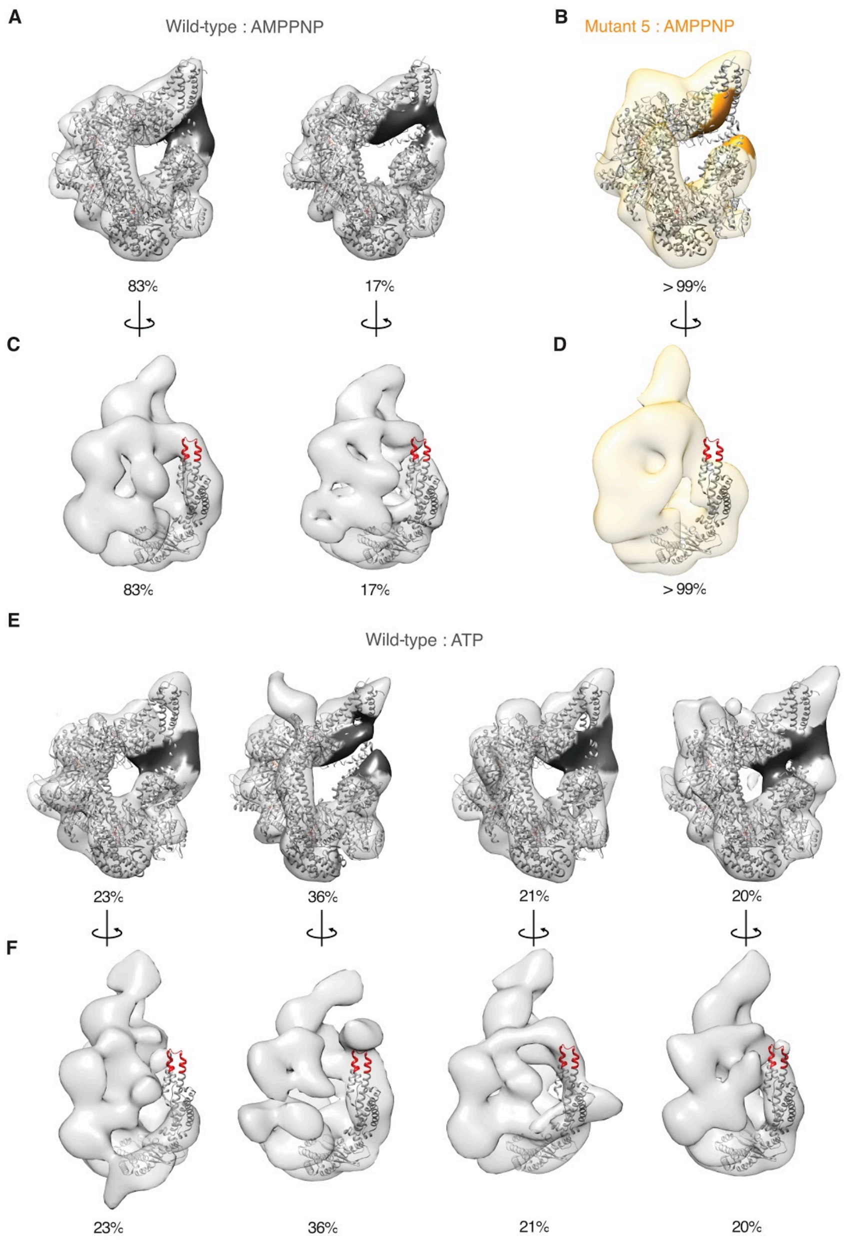


**Supplementary Figure 8 |** Negative stain reconstructions of mutant 5 and WT dynein. (A, C) Negative stain EM reconstruction of wild-type dynein (grey) in the presence of AMPPNP (EMDB: 6064 and EMDB: 6063) with the AMPPNP crystal structure (PDB: 4w8f (Bhabha *et al*, 2014)) docked-in. Major (left, EMDB: 6064) and minor (right, EMDB: 6063) conformations are shown. These data was collected in a previous study (Bhabha *et al*, 2014). (B, D) Negative stain EM density of mutant 5 (orange) in the presence of AMPPNP with the AMPPNP crystal structure (PDB: 4w8f (Bhabha *et al*, 2014)) docked-in. (A, B) Area of weak density in the AAA5 region of minor wild-type conformation (dark grey) and for mutant 5 (bright orange) are highlighted. (C, D) N-terminus of linker in crystal structure is highlighted in red and only linker and AAA1 of crystal structure are shown. (E, F) Negative stain EM density data from a previous study (Bhabha *et al*, 2014) analyzed in the light of our new findings with the AMPPNP crystal structure docked-in (PDB: 4w8f) (Bhabha *et al*, 2014). (E) Area of gap in density in the AAA5 region is highlighted in dark grey (EMDB: 6065-6068; from left to right, respectively). (F) N-terminus of linker in crystal structure is highlighted in red and only linker and AAA1 of crystal structure are shown (EMDB: 6065-6068; from left to right, respectively).

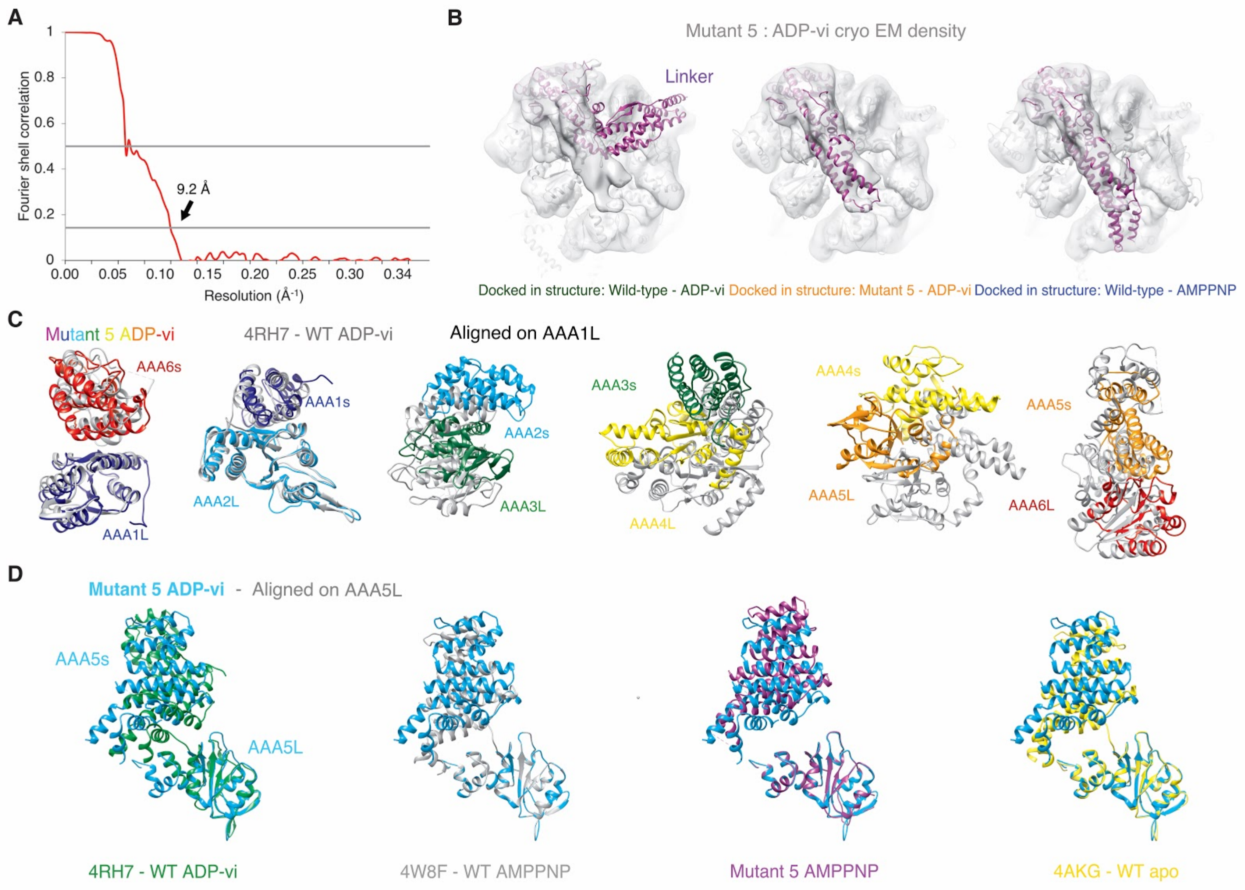


**Supplementary Figure 9 |** Cryo-EM analysis of mutant 5 in the presence of ADP-vanadate. (A) Plot of Fourier Shell Correlation (FSC) for mutant 5 in the presence of ADP-vanadate. (B) Cryo-EM reconstruction of mutant 5 with ADP-vanadate (grey) fitted with models of human cytoplasmic dynein 2 in the ADP-vi state (left - PDB: 4RH7 (Schmidt *et al*, 2015)), yeast cytoplasmic dynein mutant 5 in ADP-vi state (middle - this study), and yeast cytoplasmic dynein in the AMPPNP state (right - PDB: 4W8F (Bhabha *et al*, 2014)). For the mutant 5 ADP-vi state only the part of the linker with sufficient density was fitted. (C) Domain movements between mutant 5 and wild-type (PDB: 4RH7 (Schmidt *et al*, 2015)) in the presence of ADP-vi. The two structures were aligned on AAA1L (matchMaker in Chimera (Pettersen *et al*, 2004)). (D) Domain movements between mutant 5 in the presence of ADP-vi and - from left ro right - wild-type in the ADP-vi state (PDB: 4RH7 (Schmidt *et al*, 2015)), wild-type in the AMPPNP state (PDB: 4W8F (Bhabha *et al*, 2014)), class 1 of mutant 5 with AMPPNP, and wild-type in the apo state (PDB: 4AKG (Schmidt *et al*, 2012)). For every structure the domains are aligned on AAA5L (matchMaker in Chimera (Pettersen *et al*, 2004)).

#### Supplementary Note 1

##### The coiled-coil stalk of the dynein motor domain is strikingly conserved in length

To better understand dynein's coiled-coil stalk, we obtained a dataset of 677 unique dynein heavy chain sequences from 229 fully sequenced eukaryotic genomes including sequences from cytoplasmic, axonemal and intraflagellar transport (IFT) dyneins. Since our analysis was focused on the motor domain and in particular on the stalk, we pruned our data set (see protocol for pruning below), and aligned 534 motor domain sequences using MAFFT (Katoh *et al*, 2002; Alva *et al*, 2016).

We defined boundaries of CC1 and CC2 of the stalk using anchor residues that are very well conserved: P2989 and P3101 define CC1, and P3228 and Y3333 define CC2 (Fig. 1A, residue numbers here and throughout the manuscript correspond to yeast cytoplasmic dynein, unless otherwise specified). When we first analysed the data set (sequences we used in this study were compiled in 2014), we noticed that the length of CC1 and CC2 are extremely conserved: 501 out of 534 sequences have exactly 115 residues in CC1 and 506 out of 534 sequences have exactly 106 residues in CC2 (Supplementary Fig. 1H, I). However, since the initial analysis was carried out, several sequences have been replaced with newer corrected sequences which do not contain these insertions/deletions, suggesting that in reality closer to 1% of sequences deviate from the conserved stalk lengths (Supplementary Fig. 1J, K, Supplementary Table 1). Furthermore, we note that the length and size of other subdomains of the dynein motor domain are not as well conserved (Supplementary Fig. 1A, B). Surprisingly, despite the conservation of length, the primary sequence of the stalk is not particularly well conserved (Supplementary Fig. 1C, D). We also investigated other features such as hydrophobicity and charge variation but no clear patterns of conservation were observed in the stalk (Supplementary Fig. 1E-G).

Our initial analysis revealed a handful of sequences that varied in length from 115 residues in CC1 and 106 residues in CC2 (Supplementary Fig. 1H, I). Coiled-coils contain a repeating pattern of 7 residues (heptad repeat) consisting of charged and hydrophobic residues. The hydrophobic stretch of the heptad repeat in one coil interacts with the hydrophobic region of the heptad repeat on the other coil, thus generating a stable coiled-coil. Based on the conserved structural motifs in the coiled-coil, one may expect that, for example, a deletion of 7 residues in CC1 would correspond to a deletion of 7 residues in CC2, in order to maintain the interaction between the two coils. Indeed, previous work has shown that insertions and deletions of the same number of residues in both sides does not alternate velocity and ATPase activity significantly (Carter *et al*, 2008). Surprisingly, however, the variants in our dataset contained either insertions or deletions in one of the coils, but not in both simultaneously.

Interestingly, the sequences of all three mutants that showed diffusive movement have been updated in the databases and do not show any insertions or deletions anymore. Based on currently available sequences, most likely dyneins with this phenotype do not exist; however, we serendipitously stumbled upon these insertions and deletions in regions of the stalk, which do mediate communication between the AAA ring and MTBD, and these mutations shed light on the dynein motility mechanism.

#### Protocol for sequence alignments and design of stalk mutants

All files listed in the following format are available to download from the Supplementary Material:

--- *Sequence-alignment.fasta* ---

**Initial dataset:**

677 unique dynein heavy chain sequences (axonemal and cytoplasmic) from 229 fully sequenced eukaryotic genomes (from Christian Zmasek, Godzik lab, Burnham)

--- *All-677-heavy-chain-sequences_MAFFA-alignment.fasta* ---

**Pruning of initial dataset:**

1. To focus on the motor domain, we eliminated tail sequences and used sequences starting at E1364 (yeast numbering)
2. We removed sequences with ambiguity in sequencing reads (i.e. contains "X" in sequence)
3. We removed incomplete sequences (e.g. no amino acids in AAA6 domain)
4. We used MAFFT on the MPI Bioinformatics Toolkit server (Alva *et al*, 2016; Katoh *et al*, 2002) to realign the remaining sequences with a 'Gap open penalty' of 1.53 and an 'Offset' of 0.0

5. We removed sequences that didn't contain a functional Walker A motif in AAA1 (mutation in K1802 - yeast numbering)
6. We deleted sequences that didn't have a functional Walker B motif in AAA1 (mutation in E1849 - yeast numbering)
7. We again used MAFFT on the MPI Bioinformatics Toolkit server to realign the remaining sequences with a 'Gap open penalty' of 1.53 and an 'Offset' of 0.0

**Final dataset:**

534 unique dynein heavy chain sequences

--- *Pruned-heavy-chain-sequences_MAFFA-alignment.fasta* ---

**Division into CC1 and CC2 stalk sequences:**

1. Based on conserved residues in the multiple sequence alignment (MSA) file we used P2989 to P3103 for CC1 and P3228 to Y3333 for CC2 anchor points (yeast numbering)
1. We then truncated the final MSA file to CC1 only, to CC2 only and to CC1-MTBD-CC2 only
2. We realigned all three MSA using MAFFT on the MPI Bioinformatics Toolkit server with a 'Gap open penalty' of 1.53 and an 'Offset' of 0.0

--- *Pruned-heavy-chain-sequences_CC1_2989-3103_MAFFA-alignment.fasta* ---

--- *Pruned-heavy-chain-sequences_CC1-MTBD-CC2_2989-3333_MAFFA-alignment.fasta* ---

--- *Pruned-heavy-chain-sequences_CC2_3228-3333_MAFFA-alignment.fasta* ---

**Identification of outlier sequences:**

As can be seen in **Fig. 1** most dynein sequences of CC1 and CC2 have a well defined length of 115 and 106, respectively. Thus, we first removed all outlier sequences, these are ones that are not 115/106 amino acids long and created new alignment files for CC1 and CC2 using MAFFT on the MPI Bioinformatics Toolkit server with a 'Gap open penalty' of 1.53 and an 'Offset' of 0.0. This left us with 476 sequences. We then added one outlier sequence back at a time and created unique alignments for all possible mutations by MAFFT on the MPI Bioinformatics Toolkit server with a 'Gap open penalty' of 1.53 and an 'Offset' of 0.0.

--- *Folder: Mutants with alignments: i.e. Mutant-01_CC1.fasta* ---

**Selections of insertions / deletions to clone:**

We decided to clone mutations (insertion / deletion) that were between well defined anchor points (highly conserved residues - see Figure below (based on PDB: 4rh7) (Schmidt *et al*, 2015)), which allowed us to generate the mutant accurately in the yeast dynein background.

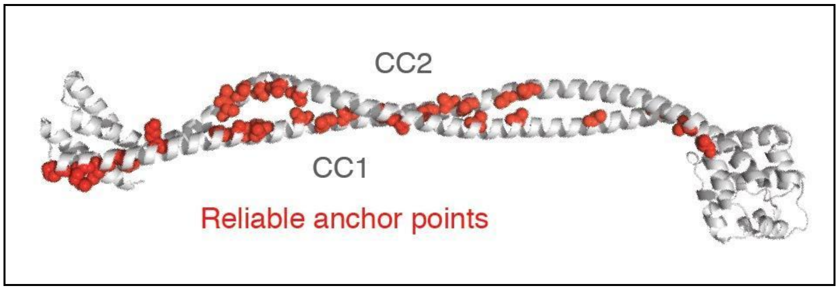


The identification of reliable anchor residues indicates regions of high sequence conservation in the stalk. In regions where reliable anchor residues were not identified, the sequence was more variable, and thus precluded us from designing meaningful insertion/deletion mutants in these regions with confidence, even though some outlier sequences did show insertions and deletions around these regions. Similar to the initial data set, our 18 mutants arise from various species and include cytoplasmic as well as axonemal dynein sequences (Supplementary Table 1).

**Updated information with newly deposited sequence data:**

We checked the sequence databases again for the sequences that have insertions or deletions and found that many were updated and corrected in the meantime (last checked on NCBI on December 22nd, 2017). This changed the percentage of outlier sequences from about five to one percent (Supplementary Fig 1H-K). We also indicated the sequences that changed in Supplementary Table 1. Note that the sequences changed on NCBI, but not necessary on EBI or uniprot.

#### Supplementary Note 2

##### Single-molecule phenotypes of stalk mutants

Using the single-molecule TIRF assay we observed a variety of phenotypes for all 18 stalk mutations (Fig. 1B, C). Two mutants did not show any protein expression, suggesting that these mutations may lead to protein misfolding and/or instability (Supplementary Fig. 4A). Five mutants bound microtubules but did not show any single-molecule movement. These mutants are clustered in the proximal region of the stalk, close to the AAA ring (Supplementary Fig. 1L), and are in regions that are important for interaction with the buttress (Kon *et al*, 2012). Based on previous structural work (Schmidt *et al*, 2015; Kon *et al*, 2012), the interaction between the stalk and buttress plays a key role in dynein motility. Therefore, one may expect mutations in the stalk region at the stalk-buttress interface to be accompanied by corresponding mutations in the buttress that have co-evolved to maintain the interaction. However, we did not identify such corresponding mutations.

Interestingly, eight mutants showed single-molecule movement with velocities and processivity that were between ~50-100% of the wild-type protein. Remarkably, some of these mutants had relatively large insertions of 5 and 9 residues or deletions of 3 or 5 residues, but did not seem impact the communication between the AAA ring and microtubule-binding domain. However, three mutations showed diffusive behavior and were subsequently analyzed in more detail.

Overall, our results show that different mutations in overlapping regions on the stalk can result in different phenotypes. For example, the locations of mutations in mutant 4 and mutant 7 overlap with mutant 5 (Supplementary Fig. 2, 3) very closely, but both these mutants behave more similarly to the wild-type protein (Supplementary Fig. 4B), with clear directional movement and only slightly decreased velocity and processivity (Supplementary Fig. 4B-E). In the case of mutant 5, mutants 4 and 7 serve as important controls, showing that we have narrowed down an exact region of the stalk that is necessary for allosteric communication. The effect of mutant 5 could be a combination of position, exact length of deletion, and change in structural properties of the new stalk configuration. However, mutants 4 and 7 illustrate that insertions and deletions in overlapping areas can have very small effects on motility.

Similarly, mutants with insertions and deletions containing the same number of residues in the same coiled-coil can move very differently, suggesting that it is not simply the total length of the coil that dictates function. For instance, mutant 1 and mutant 18 have an insertion of 1 residue in CC1, but mutant 1 motility is very similar to wild-type motility, whereas mutant 18 does not move at all. For the three mutants that have an insertion of 1 residue in CC2, mutant 2 and mutant 13 show diffusive like behavior while mutant 15 was destabilized such that it did not express at all.

#### Supplementary Note 3

##### Negative stain electron microscopy of mutant 5

Prior to the Cryo-EM study we used a negative stain EM based assay with the goal of assessing the linker conformation and/or flexibility. Due to the large-scale conformational change in the linker, we were previously able to identify distinct linker conformations using 3D classification and refinement from negative stain EM data (Bhabha *et al*, 2014). For this assay, we used the ATP analog AMPPNP to mimic an ATP-bound state of the enzyme at AAA1 and AAA3. In this state, the linker in the wild-type enzyme is straight and docked to the AAA ring at AAA5 (Bhabha *et al*, 2014). 3D classification and refinement of negative stain EM data of mutant 5 showed weaker density for the N-terminal region of the linker, suggesting that the position of the linker in mutant 5 may be more flexible than in wild-type (Supplementary Fig. 8 A-D). More strikingly, however, we observed a "gap" (missing density) in the ring at the region of AAA5 (Supplementary Fig. 8B), in comparison to a more closed state of the wild-type motor (Bhabha *et al*, 2014).

#### Supplementary Tables

| Mutation Number / dimeric or monomeric | Strain name | Organism | NCBI - proteinBLAST | EBI - European Bioinformatics Institute | Uniprot - HMMER search |
|---|---|---|---|---|---|
| WT / dimer | VY208 | Saccharomyces cerevisiae | https://www.ncbi.nlm.nih.gov/protein/767040268?report=genbank&log$=protalign&blast_rank=2&RID=XUTTCCYZ014 | - | http://www.uniprot.org/uniprot/P36022 |
| 1 / dimer | VY1044 | Gorilla gorilla gorilla | UPDATED: https://www.ncbi.nlm.nih.gov/protein/XP_004044002 | https://www.ebi.ac.uk/ebisearch/search.ebi?db=allebi&query=ENSGGOP00000020228 | http://www.uniprot.org/uniprot/G3RWP4 |
| 2 / dimer | VY1045 | Helobdella robusta | UPDATED: https://www.ncbi.nlm.nih.gov/protein/675890198?report=genbank&log$=protalign&blast_rank=1&RID=XUU15W9Y014 | - | http://www.uniprot.org/uniprot/T1G9C1 |
| 3 / dimer | VY1046 | Takifugu rubripes | UPDATED: https://www.ncbi.nlm.nih.gov/protein/XP_011616710?report=genbank&log$=protalign&blast_rank=1&RID=41WVKX4Y014 | https://www.ebi.ac.uk/ebisearch/search.ebi?db=allebi&query=ENSTRUP00000031696 | http://www.uniprot.org/uniprot/H2U434 |
| 4 / dimer | VY1047 | Branchiostoma floridae | UPDATED (Branchiostoma belcheri): https://www.ncbi.nlm.nih.gov/protein/XP_019639192?report=genbank&log$=protalign&blast_rank=1&RID=41WVXYCX014 | - | - |
| 5 / dimer | VY1048 | Nasonia vitripennis | UPDATED: https://www.ncbi.nlm.nih.gov/protein/XP_008209982?report=genbank&log$=protalign&blast_rank=1&RID=41WWA4A8014 | - | http://www.uniprot.org/uniprot/K7J523 |

| | | | | | |
|---|---|---|---|---|---|
| 6 / dimer | VY1049 | Takifugu rubripes | - | https://www.ebi.ac.uk/ebisearch/search.ebi?db=allebi&query=ENSTRUP00000000144 | http://www.uniprot.org/uniprot/H2RJ31 |
| 7 / dimer | VY1050 | Cavia porcellus | UPDATED: https://www.ncbi.nlm.nih.gov/protein/XP_003463142?report=genbank&log$=protalign&blast_rank=2&RID=41Y4RCPD015 | https://www.ebi.ac.uk/ebisearch/search.ebi?db=allebi&query=ENSCPOP00000003676 | http://www.uniprot.org/uniprot/H0V2C0 |
| 8 / dimer | VY1051 | Takifugu rubripes | UPDATED: https://www.ncbi.nlm.nih.gov/protein/XP_003966059?report=genbank&log$=protalign&blast_rank=1&RID=41WXNEUS015 | https://www.ebi.ac.uk/ebisearch/search.ebi?db=allebi&query=ENSTRUP00000008414 | http://www.uniprot.org/uniprot/H2S7P2 |
| 9 / dimer | VY1052 | Takifugu rubripes | UPDATED: https://www.ncbi.nlm.nih.gov/protein/XP_003966059?report=genbank&log$=protalign&blast_rank=1&RID=41WY1J2K014 | https://www.ebi.ac.uk/ebisearch/search.ebi?db=allebi&query=ENSTRUP00000008415 | http://www.uniprot.org/uniprot/H2S7P3 |
| 10 / dimer | VY1053 | Ciona intestinalis | UPDATED: https://www.ncbi.nlm.nih.gov/protein/XP_018671050?report=genbank&log$=protalign&blast_rank=1&RID=41WYCT79014 | https://www.ebi.ac.uk/ebisearch/search.ebi?db=allebi&query=ENSCINP00000008812 | - |
| 11 / dimer | VY1054 | Ciona savignyi | UPDATED (Ciona intestinalis): https://www.ncbi.nlm.nih.gov/protein/XP_009858173?report=genbank&log$=protalign&blast_rank=1&RID=41WYPZH4015 | https://www.ebi.ac.uk/ebisearch/search.ebi?db=allebi&query=ENSCSAVP00000008997 | http://www.uniprot.org/uniprot/H2YUI7 |
| 12 / dimer | VY1062 | Ciona savignyi | UPDATED (Ciona intestinalis): https://www.ncbi.nlm.nih.gov/protein/XP_009858173?report=genbank&log$=protalign&blast_rank=1&RID=41WZ437X015 | https://www.ebi.ac.uk/ebisearch/search.ebi?db=allebi&query=ENSCSAVP00000009000 | http://www.uniprot.org/uniprot/H2YUJ0 |
| 13 / dimer | VY1056 | Ciona savignyi | UPDATED (Ciona intestinalis): https://www.ncbi.nlm.nih.gov/protein/XP_018669141?report=genbank&log$=protalign&blast_rank=1&RID=41WZF9S8014 | https://www.ebi.ac.uk/ebisearch/search.ebi?db=allebi&query=ENSCSAVP00000010325 | http://www.uniprot.org/uniprot/H2YYB4 |

| | | | | | |
|---|---|---|---|---|---|
| 14 / dimer | VY1057 | Ciona intestinalis | - | https://www.ebi.ac.uk/ebisearch/search.ebi?db=allebi&query=ENSCINP00000011393 | - |
| 15 / dimer | VY1058 | Anolis carolinensis | UPDATED: https://www.ncbi.nlm.nih.gov/protein/XP_003217173?report=genbank&log$=protalign&blast_rank=1&RID=41X056ZX015 | https://www.ebi.ac.uk/ebisearch/search.ebi?db=allebi&query=ENSACAP00000016375 | http://www.uniprot.org/uniprot/G1KSW2 |
| 16 / dimer | VY1059 | Gallus gallus | UPDATED: https://www.ncbi.nlm.nih.gov/protein/XP_015137732.1?report=genbank&log$=prottop&blast_rank=8&RID=41X0FWDX015 | - | - |
| 17 / dimer | VY1060 | E. cuniculi | https://www.ncbi.nlm.nih.gov/protein/19074673?report=genbank&log$=protalign&blast_rank=1&RID=XWEHW0TG015 | - | http://www.uniprot.org/uniprot/Q8SR52 |
| 18 / dimer | VY1061 | Ciona intestinalis | - | https://www.ebi.ac.uk/ebisearch/search.ebi?db=allebi&query=ENSCINP00000011395 | - |
| WT / monomer | VY137 | See WT / dimer | | | |
| 2 / monomer | VY1063 | See 2 / dimer | | | |
| 5 / monomer | VY1065 | See 5 / dimer | | | |
| 13 / monomer | VY1071 | See 13 / dimer | | | |

**Supplementary Table 1 |** Annotation of all dynein stalk mutant strains used in this study. The VY208 genotype is: MATa; his3-11,15; ura3-1; leu2-3,112; ade2-1; trp1-1; PEP4::HIS5; PRB1D pDyn-pGAL-ZZ-TEV-GFP-3XHA-GST-D6-DYN1-gsDHA:Kan) and the VY137 genotype is: PGal:ZZ:Tev:GFP:HA:D6 MATa; his3-11,15; ura3-1; leu2-3,112; ade2-1; trp1-1; PEP4::HIS5; PRB1D. All sequences that say “UPDATED” do not have any insertions or deletions anymore (based on NCBI (December 22nd, 2017)). For more details see Supplementary Note 1.

“-” indicates that the sequence was not found (sequence identity less than 60%).

| Construct | Nucleotide | $K_d$ [MT] | $B_M$ | $k_{basal}$ |
|---|---|---|---|---|
| Wild-type | ATP | 5.22 ± 0.92 $\mu$M | 0.23 ± 0.17 | 0.02 ± 0.01 |
| Wild-type | apo | 0.78 ± 0.27 $\mu$M | 0.86 ± 0.02 | 0.02 ± 0.01 |
| Wild-type | AMPPNP | 1.22 ± 0.72 $\mu$M | 0.90 ± 0.04 | 0.03 ± 0.01 |
| Mutant 2 | ATP | 2.62 ± 0.89 $\mu$M | 0.15 ± 0.12 | 0.01 ± 0.01 |
| Mutant 2 | apo | 5.83 ± 0.04 $\mu$M | 0.32 ± 0.01 | 0.01 ± 0.00 |
| Mutant 2 | AMPPNP | 3.93 ± 1.59 $\mu$M | 0.31 ± 0.05 | 0.03 ± 0.01 |
| Mutant 5 | ATP | 4.10 ± 1.28 $\mu$M | 0.26 ± 0.01 | 0.05 ± 0.00 |
| Mutant 5 | apo | 5.89 ± 1.38 $\mu$M | 0.25 ± 0.05 | 0.02 ± 0.01 |
| Mutant 5 | AMPPNP | 3.08 ± 2.06 $\mu$M | 0.20 ± 0.06 | 0.02 ± 0.01 |
| Mutant 13 | ATP | 4.00 ± 0.79 $\mu$M | 0.28 ± 0.01 | 0.02 ± 0.00 |
| Mutant 13 | apo | 6.19 ± 0.99 $\mu$M | 0.33 ± 0.06 | 0.02 ± 0.00 |
| Mutant 13 | AMPPNP | 3.66 ± 0.08 $\mu$M | 0.25 ± 0.02 | 0.02 ± 0.00 |

**Supplementary Table 2 |** Microtubule affinity measurements. The data were fit to the following equation $k_{obs} = (B_M - k_{basal}) \frac{[MT]}{K_d + [MT]} + k_{basal}$ ($B_M$ maximum binding, $K_d$ dissociation constant). Values are shown as averages of triplicates ± standard deviation.

| **Mutation** | **Organism** | **$K_M$ [MT]** | **$k_{cat}$** | **$k_{basal}$** | **Reference** |
|---|---|---|---|---|---|
| Wild-type | Yeast | 0.59 ± 0.28 $\mu$M | 14.1 ± 0.36 $s^{-1}$ | 3.74 ± 0.35 $s^{-1}$ | Cho et al. JCB 2008 (Cho *et al*, 2008) |
| AAA3 (E2488Q) | Yeast | 0.03 ± 0.01 $\mu$M | 1.38 ± 0.14 $s^{-1}$ | 0.30 ± 0.05 $s^{-1}$ | Cho et al. JCB 2008 |
| AAA4 (E2819Q) | Yeast | 0.09 ± 0.03 $\mu$M | 10.6 ± 0.72 $s^{-1}$ | 3.36 ± 0.59 $s^{-1}$ | Cho et al. JCB 2008 |
| Wild-type | Yeast | - | 20 ± 4 $s^{-1}$ | 6 ± 2 $s^{-1}$ | Carter et al. Science 2008 (Carter *et al*, 2011) |
| Removal of 7 heptads in stalk | Yeast | - | 21 ± 2 $s^{-1}$ | 13 ± 2 $s^{-1}$ | Carter et al. Science 2008 |
| Insertion of 7 heptads in stalk | Yeast | - | 21 ± 5 $s^{-1}$ | 6 ± 2 $s^{-1}$ | Carter et al. Science 2008 |
| Wild-type | Yeast | 1.06 ± 0.16 $\mu$M | 16.75 ± 0.49 $s^{-1}$ | 3.51 ± 0.31 $s^{-1}$ | Toropova et al. eLife 2014 (Toropova *et al*, 2014) |
| AAA1 | Yeast | - | - | ~ 1 $s^{-1}$ | Toropova et al. eLife 2014 |
| AAA5 - linker docking (F3446D, R3445E, K3438E) | Yeast | - | - | ~ 2 $s^{-1}$ | Toropova et al. eLife 2014 |
| Wild-type | D.discoideum | 33.3 ± 2.6 $\mu$M | 105.2 ± 4.2 $s^{-1}$ | 8.7 ± 0.8 $s^{-1}$ | Kon et al. NSMB 2009 (Kon *et al*, 2009) |
| Fixed $\alpha$ registry (oxidized) | D.discoideum | 5.0 ± 1.0 $\mu$M | 158.0 ± 1.7 $s^{-1}$ | 128.4 ± 6.6 $s^{-1}$ | Kon et al. NSMB 2009 |
| Fixed $\beta$ + registry (oxidized) | D.discoideum | 19.2 ± 1.7 $\mu$M | 17.0 ± 0.6 $s^{-1}$ | 3.3 ± 0.1 $s^{-1}$ | Kon et al. NSMB 2009 |

| | | | | | |
|---|---|---|---|---|---|
| Fixed $\beta$ - registry (oxidized) | D.discoideum | 20.4 ± 4.6 $\mu$M | 112.9 ± 4.1 $s^{-1}$ | 74.0 ± 2.1 $s^{-1}$ | Kon et al. NSMB 2009 |
| Delta buttress (Q3824-E3864) | D.discoideum | - | - | ~ 90 $s^{-1}$ | Kon et al. Nature 2012 (Kon *et al*, 2012) |
| Delta c-terminus (S4416-I4730) | D.discoideum | - | - | ~ 10 $s^{-1}$ | Kon et al. Nature 2012 |
| Wild-type | Yeast | - | ~ 17 $s^{-1}$ | ~ 3 $s^{-1}$ | Bhabha et al. Cell 2014 (Bhabha *et al*, 2014) |
| AAA2 - linker docking (A2121G, T2122G, L2123G) | Yeast | - | ~ 7 $s^{-1}$ | ~ 3 $s^{-1}$ | Bhabha et al. Cell 2014 |
| AAA2 - linker docking (R2183A) | Yeast | - | ~ 5 $s^{-1}$ | ~ 2 $s^{-1}$ | Bhabha et al. Cell 2014 |
| GST-Dynein (1219-4093) | Yeast | 0.39 ± 0.06 $\mu$M | 16.1 ± 0.3 $s^{-1}$ | - | Reck-Peterson et al. Cell 2006 (Reck-Peterson *et al*, 2006) |
| Dynein-GST (1219-4093) | Yeast | - | 4.3 ± 0.3 $s^{-1}$ | - | Reck-Peterson et al. Cell 2006 |
| GST-Dynein (1390-4093) | Yeast | - | - | ~ 1 $s^{-1}$ | Reck-Peterson et al. Cell 2006 |

**Supplementary Table 3 |** ATPase rates for dynein mutations in the literature.

| Construct | $K_M$ [MT] | $k_{cat}$ | $k_{basal}$ |
|---|---|---|---|
| Wild-type | 0.50 ± 0.17 $\mu$M | 15.18 ± 1.18 $s^{-1}$ | 0.75 ± 0.34 $s^{-1}$ |
| Mutant 2 | - | - | 6.23 ± 2.25 $s^{-1}$ |
| Mutant 5 | - | - | 13.80 ± 0.50 $s^{-1}$ |
| Mutant 13 | - | - | 14.95 ± 0.35 $s^{-1}$ |

**Supplementary Table 4 |** ATPase assay rate measurements. The data were fit to the following equation $k_{obs} = (k_{cat} - k_{basal}) \frac{[MT]}{K_M + [MT]} + k_{basal}$. Values are shown as averages of triplicates ± standard deviation. - not measurable

| **Data Collection (Cryo-EM)** | **Mutant 5 + AMPPNP** | | **Mutant 5 + ATP-vi** | |
|---|---|---|---|---|
| Microscope | Titan Krios | | Arctica | |
| Camera | K2 | | K2 | |
| Magnification | 22,500 | | 36,000 | |
| Voltage (kV) | 300 | | 200 | |
| Electron dose (e-/pixel/second) | 10 | | 8 | |
| Focus range (μm) | 1.5-3.0 | | 1.5-3.0 | |
| Pixel size (Å) | 1.31 | | 1.156 | |
| Number of images/movies | 1200 | | 664 | |
| **Reconstruction** | | | | |
| Particles selected after 2D classification (no.) | 310,085 | | 35,565 | |
| CTF correction tool | GCTF 1.0.6 | | ctffind 4.1.10 | |
| Particle picking method | Gaussian blobs | | Gaussian blobs | |
| Ab-initio models generated (no.) | 4 | | 5 | |
| Last round of 3D heterogeneous refinement (no.) | 4 | | 5 | |
| Class name | **Class 1** | **Class 2** | **Class 1** | **Class 2 (not shown)** |
| Point group symmetry | C1 | C1 | C1 | C1 |
| Final particles (no.) | 97,008 | 39,048 | 8,653 | 6,629 |
| Resolution (Å) | 7.7 | 7.6 | 9.2 | 16.6 |
| B-factor ($Å^2$) | -400 | -400 | -400 | -400 |
| Modelling from 4W8F domains | rigid-body (Chimera) | rigid-body (Chimera) + refinement (PHENIX) | rigid-body (Chimera) | None |

**Supplementary Table 5 |** Statistics on cryo-EM data collection and processing.

#### Supplementary Movies

Supplementary-Movie_1: **Single-molecule movement of wild-type dynein with 1 mM ATP**

The movie shows single molecules of Halo488-labeled wild-type dynein (green) bound to surface-immobilized microtubules (blue) in the presence of 1 mM ATP. Time is in min:sec.

Supplementary-Movie_2: **Single-molecule movement of mutant 2 dynein with 1 mM ATP**

The movie shows single molecules of Halo488-labeled mutant 2 dynein (green) bound to surface-immobilized microtubules (blue) in the presence of 1 mM ATP. Time is in min:sec.

Supplementary-Movie_3: **Single-molecule movement of mutant 5 dynein with 1 mM ATP**

The movie shows single molecules of Halo488-labeled mutant 5 dynein (green) bound to surface-immobilized microtubules (blue) in the presence of 1 mM ATP. Time is in min:sec.

Supplementary-Movie_4: **Single-molecule movement of mutant 13 dynein with 1 mM ATP**

The movie shows single molecules of Halo488-labeled mutant 13 dynein (green) bound to surface-immobilized microtubules (blue) in the presence of 1 mM ATP. Time is in min:sec.

Supplementary-Movie_5: **Single-molecule movement of wild-type dynein without ATP**

The movie shows single molecules of Halo488-labeled wild-type dynein (green) bound to surface-immobilized microtubules (blue) without ATP. Time is in min:sec.

Supplementary-Movie_6: **Single-molecule movement of mutant 2 dynein without ATP**

The movie shows single molecules of Halo488-labeled mutant 2 dynein (green) bound to surface-immobilized microtubules (blue) without ATP. Time is in min:sec.

Supplementary-Movie_7: **Single-molecule movement of mutant 5 dynein without ATP**

The movie shows single molecules of Halo488-labeled mutant 5 dynein (green) bound to surface-immobilized microtubules (blue) without ATP. Time is in min:sec.

Supplementary-Movie_8: **Single-molecule movement of mutant 13 dynein without ATP**

The movie shows single molecules of Halo488-labeled mutant 13 dynein (green) bound to surface-immobilized microtubules (blue) without ATP. Time is in min:sec.

Supplementary-Movie_9: **Gilding of microtubules by wild-type dynein**

The movie shows the gliding of microtubules (green) by surface-immobilized wild-type dynein in the presence of 1 mM ATP. Time is in min:sec. Note: Even though Kinesin is present in the assay it is not shown for clarity reasons.

Supplementary-Movie_10: **Gilding of microtubules by dynein with stalk mutation 2**

The movie shows the gliding of microtubules (green) by surface-immobilized mutant 2 dynein in the presence of 1 mM ATP. Time is in min:sec. Note: Even though Kinesin is present in the assay it is not shown for clarity reasons.

Supplementary-Movie_11: **Gilding of microtubules by dynein with stalk mutation 5**

The movie shows the gliding of microtubules (green) by surface-immobilized mutant 5 dynein in the presence of 1 mM ATP. Time is in min:sec. Note: Even though Kinesin is present in the assay it is not shown for clarity reasons.

Supplementary-Movie_12: **Gilding of microtubules by dynein with stalk mutation 13**

The movie shows the gliding of microtubules (green) by surface-immobilized mutant 13 dynein in the presence of 1 mM ATP. Time is in min:sec. Note: Even though Kinesin is present in the assay it is not shown for clarity reasons.

Supplementary-Movie_13: **Conformational changes in the AAA ring from mutant5 ADP-vi to mutant5 AMPPNP class 1 compared to WT ADP-vi to WT AMPPNP aligned on AAA1L.**

Morphs were generated in Chimera (matchMaker in Chimera (Pettersen *et al*, 2004)) using the mutant 5 ADP-vanadate structure and the mutant 5 AMPPNP class 1 structure as well as the ADP-vanadate bound human cytoplasmic dynein 2 structure (PDB: 4RH7) (Schmidt *et al*, 2015) and the yeast AMPPNP state structure (PDB: 4W8F (Bhabha *et al*, 2014)). Both ADP-vanadate structures were aligned to the respective AMPPNP structure (wild-type and mutant 5) using AAA1L as anchor.

Supplementary-Movie_14: **Conformational changes in the AAA ring from mutant5 ADP-vi to mutant5 AMPPNP class 1 compared to WT ADP-vi to WT AMPPNP aligned on AAA2L.**

Morphs were generated in Chimera (matchMaker in Chimera (Pettersen *et al*, 2004)) using the mutant 5 ADP-vanadate structure and the mutant 5 AMPPNP class 1 structure as well as the ADP-vanadate bound human cytoplasmic dynein 2 structure (PDB: 4RH7) (Schmidt *et al*, 2015) and the yeast AMPPNP state structure (PDB: 4W8F (Bhabha *et al*, 2014)). Both ADP-vanadate structures were aligned to the respective AMPPNP structure (wild-type and mutant 5) using AAA2L as anchor.
